# The pathogenic T42A mutation in SHP2 rewires the interaction specificity of its N-terminal regulatory domain

**DOI:** 10.1101/2023.07.10.548257

**Authors:** Anne E. van Vlimmeren, Rashmi Voleti, Cassandra A. Chartier, Ziyuan Jiang, Deepti Karandur, Preston A. Humphries, Wan-Lin Lo, Neel H. Shah

**Author notes:** These authors contributed equally (Anne E. van Vlimmeren, Rashmi Voleti).

## Abstract

Mutations in the tyrosine phosphatase SHP2 are associated with a variety of human diseases. Most mutations in SHP2 increase its basal catalytic activity by disrupting auto-inhibitory interactions between its phosphatase domain and N-terminal SH2 (phosphotyrosine recognition) domain. By contrast, some disease-associated mutations located in the ligand-binding pockets of the N- or C-terminal SH2 domains do not increase basal activity and likely exert their pathogenicity through alternative mechanisms. We lack a molecular understanding of how these SH2 mutations impact SHP2 structure, activity, and signaling. Here, we characterize five SHP2 SH2 domain ligand-binding pocket mutants through a combination of high-throughput biochemical screens, biophysical and biochemical measurements, and molecular dynamics simulations. We show that, while some of these mutations alter binding affinity to phosphorylation sites, the T42A mutation in the N-SH2 domain is unique in that it also substantially alters ligand-binding specificity, despite being 8-10 Å from the specificity-determining region of the SH2 domain. This mutation exerts its effect on sequence specificity by remodeling the phosphotyrosine binding pocket, altering the mode of engagement of both the phosphotyrosine and surrounding residues on the ligand. The functional consequence of this altered specificity is that the T42A mutant has biased sensitivity toward a subset of activating ligands and enhances downstream signaling. Our study highlights an example of a nuanced mechanism of action for a disease-associated mutation, characterized by a change in protein-protein interaction specificity that alters enzyme activation.

**Significance Statement:** The protein tyrosine phosphatase SHP2 is mutated in a variety of human diseases, including several cancers and developmental disorders. Most mutations in SHP2 hyperactivate the enzyme by destabilizing its auto-inhibited state, but several disease-associated mutations do not conform to this mechanism. We show that one such mutation, T42A, alters the ligand binding specificity of the N-terminal regulatory domain of SHP2, causing the mutant phosphatase to be more readily activated by certain upstream signals than the wild-type phosphatase. Our findings reveal a novel mode of SHP2 dysregulation that will improve our understanding of pathogenic signaling. Our study also illustrates how mutations distal to the specificity-determining region of a protein can alter ligand binding specificity.

## Introduction

SHP2 (Src homology-2 domain-containing protein tyrosine phosphatase-2) is a ubiquitously expressed protein tyrosine phosphatase, encoded by the *PTPN11* gene. It has critical roles in many biological processes, including cell proliferation, development, immune-regulation, metabolism, and differentiation (1–3). Germline *PTPN11* mutations underlie approximately 50% of all cases of Noonan Syndrome, a congenital disorder which is characterized by a wide range of developmental defects (4–6). In addition, somatic mutations in *PTPN11* are found in roughly 35% of patients with juvenile myelomonocytic leukemia (JMML), a rare pediatric cancer (7, 8). *PTPN11* mutations also occur in other types of leukemia, such as acute myeloid leukemia, acute lymphoid leukemia and chronic myelomonocytic leukemia, as well as solid tumors, albeit at lower incidence (9).

SHP2 has three globular domains: a protein tyrosine phosphatase (PTP) domain, which catalyzes the dephosphorylation of tyrosine-phosphorylated proteins, and two Src Homology 2 (SH2) domains, which are phosphotyrosine (pTyr)-recognition domains (**Figure 1A**). The SH2 domains regulate SHP2 activity by dictating localization and through allosteric control of catalytic activity. Interactions between the N-SH2 domain and PTP domain limit substrate access by blocking the catalytic site, leading to an auto-inhibited state with low basal catalytic activity. Conformational changes of the N-SH2 domain, caused by its binding to tyrosine-phosphorylated proteins, disrupt the N-SH2/PTP interaction to activate SHP2 in a ligand-dependent manner (**Figure 1B,C**) (10). Thus, the N-SH2 domain couples the localization of SHP2 to its activation by specific upstream signals.

**Figure 1.**
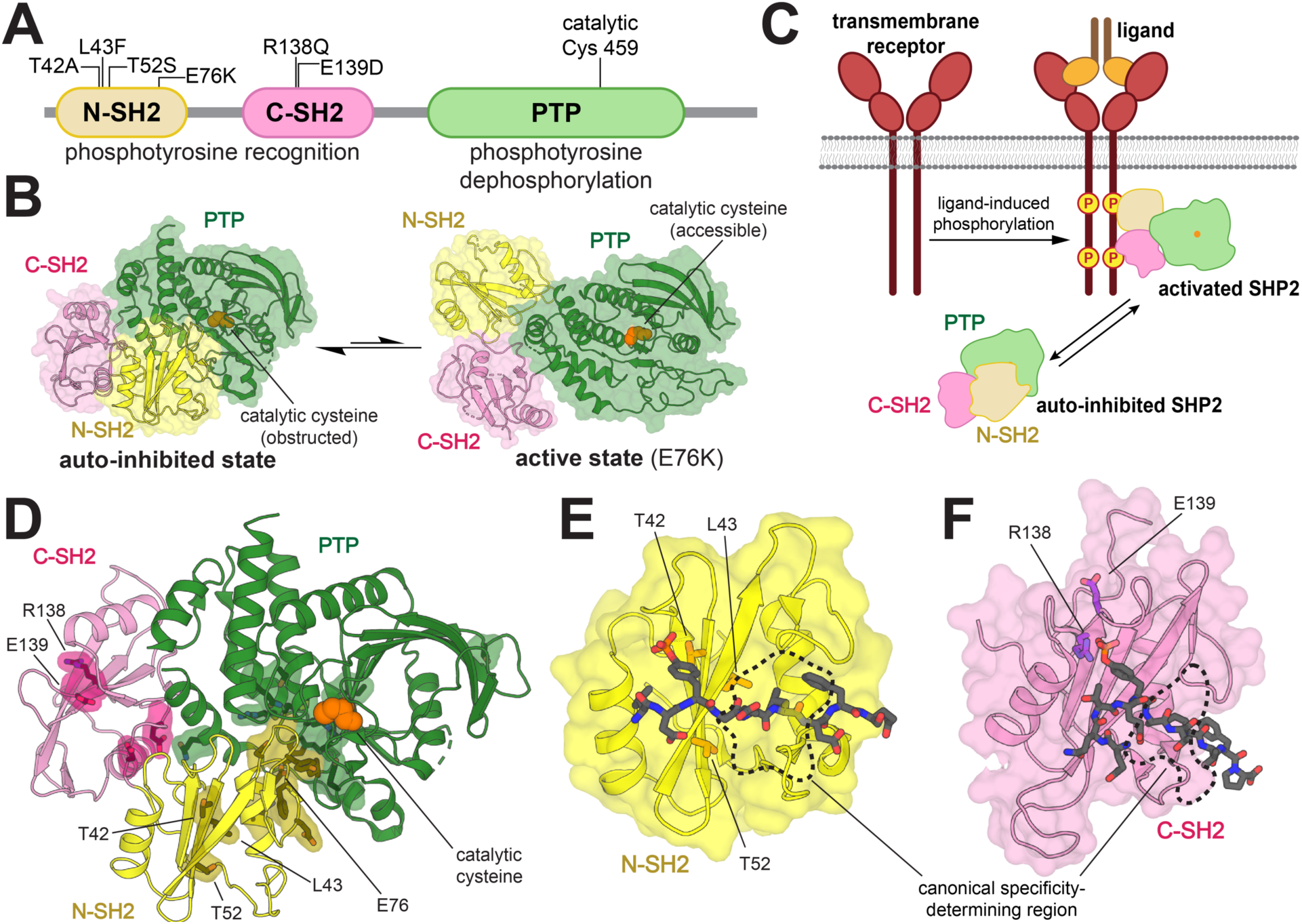
Structure and regulation of SHP2. (**A**) Domain architecture diagram of SHP2. Relevant mutations and the catalytic cysteine (Cys^459^) are indicated. (**B**) Auto-inhibited state of SHP2 (*left*, PDB: 4DGP) and representative active state of SHP2 (*right*, E76K mutant, PDB: 6CRF). (**C**) The SH2 domains of SHP2 bind to upstream phosphoproteins, inducing a conformational change that activates SHP2. (**D**) Rendering of auto-inhibited SHP2, highlighting several disease-associated mutation sites within (e.g. E76) and outside (e.g. T42) of the N-SH2/PTP interface (PDB: 4DGP). (**E**) Mutation sites in or near the N-SH2 binding pocket (PDB: 6ROY). (**F**) Mutation sites in or near the C-SH2 binding pocket (PDB: 6R5G). The primary specificity-determining regions of the SH2 domains, which dictate +1 to +5 residue preferences, are marked with black dashed lines.

Over 100 disease-associated mutations in *PTPN11* have been reported, yielding amino acid substitutions at more than 30 different residues spanning all three globular domains (4). Most known mutations in SHP2 are at the N-SH2/PTP auto-inhibitory interface and shift the conformational equilibrium of SHP2 towards the active state (**Figure 1D**) (10–13). These mutations cause SHP2 to populate a catalytically active state, largely irrespective of localization or activating stimuli. By contrast, some pathogenic mutations are found in the pTyr-binding pockets of the N- and C-SH2 domains and are mechanistically distinct, as they have the potential to change the nature of SH2-phosphoprotein interactions. T42A, a Noonan Syndrome mutation in the pTyr-binding pocket of the N-SH2 domain, has been reported to enhance binding affinity for various SHP2 interactors (**Figure 1E**) (14). This mutation is thought to make SHP2 more readily activated by upstream phosphoproteins. However, SHP2 still requires binding and localization to those phosphoproteins for functional signaling. Beyond its known effect on ligand binding affinity, the precise effect of the T42A mutation on specific cell signaling processes remains elusive. Two nearby mutations in the N-SH2 domain, L43F and T52S (**Figure 1E**), are associated with non-syndromic heart defects and JMML, respectively, but very little is known about their effects on SHP2 activity (15, 16). The C-SH2 domain mutant R138Q has been identified in melanoma, whereas the E139D mutation is associated with both JMML and Noonan Syndrome (**Figure 1F**) (9, 17). Insights into the molecular mechanisms underlying these pathogenic pTyr-binding pocket mutations could further our understanding of how they dysregulate cellular signaling, and in turn, cause tumorigenesis or aberrant development.

In this study, we extensively characterize the binding properties of five disease-associated SH2-domain mutations in SHP2. Through a series of biophysical measurements and high-throughput peptide-binding screens, we demonstrate that the T42A mutation in the N-SH2 domain is unique among these mutations in that it changes the sequence specificity of the N-SH2 domain. Notably, the T42A mutation does not lie in a canonical specificity determining region for SH2 domains (**Figure 1E, F**) (18, 19). Through molecular dynamics simulations and further biochemical experiments, we identify structural changes caused by the T42A mutation that explain its altered ligand binding specificity. We show that this change in specificity within the N-SH2 domain results in sequence-dependent changes to the activation of full-length SHP2 by phosphopeptide ligands. Finally, we demonstrate that these findings are robust in a cellular context, by showing that SHP2^T42A^ binds tighter than SHP2^WT^ to several full-length phosphoprotein interactors and enhances downstream signaling. Our results suggest that the pathogenicity of SHP2^T42A^ could be due to biased sensitization to specific upstream signaling partners, caused by rewiring of the interaction specificity of the N-SH2 domain.

## Results

### Mutations in the SH2 domains of SHP2 impact both binding affinity and sequence specificity

Mutations in the SH2 domains of SHP2 potentially change both the affinity and the specificity of the SH2 domain, thereby affecting SH2 domain functions such as recruitment, localization, and allosteric regulation of SHP2 activity. We focused on three mutations in the N-SH2 domain (T42A, L43F, and T52S) that are disease-relevant and close to the pTyr-binding pocket (**Figure 1E**). These mutations do not cause a substantial increase in basal phosphatase activity, in contrast to the E76K mutation, which lies at the auto-inhibitory interface and strongly enhances phosphatase activity (**Figure S1A and Table S1A**) (17, 20, 21). We also studied R138Q and E139D, two disease-associated mutations in the pTyr-binding pocket of the C-SH2 domain (**Figure 1F**). E139D causes a 15-fold increase in basal phosphatase activity (**Figure S1A and Table S1**), as has been reported previously (17, 22). The R138Q mutation is expected to disrupt phosphopeptide binding, as Arg^138^ directly coordinates the phosphoryl group of phosphotyrosine ligand (23, 24). This mutation had no impact on the basal catalytic activity of SHP2 (**Figure S1A and Table S1**). We also measured the melting temperatures of these mutants, as a proxy for the conformational state of the protein (25). Only the L43F mutant showed a modest decrease in melting temperature, suggestive of a slightly more open conformation than wild-type (**Figure S1B**). By comparison, the E76K mutant had a much lower melting temperature, consistent with its open conformation.

Using a fluorescence polarization assay, we measured the binding affinity of a fluorescent phosphopeptide derived from a known SHP2 binding site (pTyr^1179^) on insulin receptor substrate 1 (IRS-1) against all the four N-SH2 domain variants (**Figure S2A and Table S2**) (26). We found that N-SH2^T42A^ binds 5-fold tighter to this peptide compared to N-SH2^WT^, consistent with previous literature demonstrating enhanced binding for N-SH2^T42A^ (14). Next, in competition binding experiments, we tested 8 unlabeled phosphopeptides derived from known SHP2 binders, and one unlabeled phosphopeptide (Imhof-9) based on an unnatural ligand discovered in a previously reported peptide screen (**Figure 2, Figure S2B, and Table S2**) (26–32). We observed a broad range of effects on binding affinity for N-SH2^T42A^. Compared to N-SH2^WT^, N-SH2^T42A^ displayed a 28-fold increase in affinity for the PD-1 pTyr^223^ phosphopeptide, while a 20-fold increase was observed for Gab2 pTyr^614^ (**Figure 2B**). The increase in affinity for other peptides was more moderate, ranging from 4- to 6-fold. This suggests that the T42A mutation selectively enhances the affinity of the N-SH2 domain for specific peptides. By contrast, for N-SH2^L43F^ and N-SH2^T52S^, we observed a 2- to 3-fold increase in binding affinity for some peptides when compared to N-SH2^WT^ (**Figure 2C,D**). To confirm that the effects were specific to these ligand-binding pocket mutations, we measured the binding affinity of N-SH2^E76K^ to four peptides. All N-SH2^E76K^ affinities were similar to those for N-SH2^WT^ (**Figure S2C, Table S2**).

**Figure 2.**
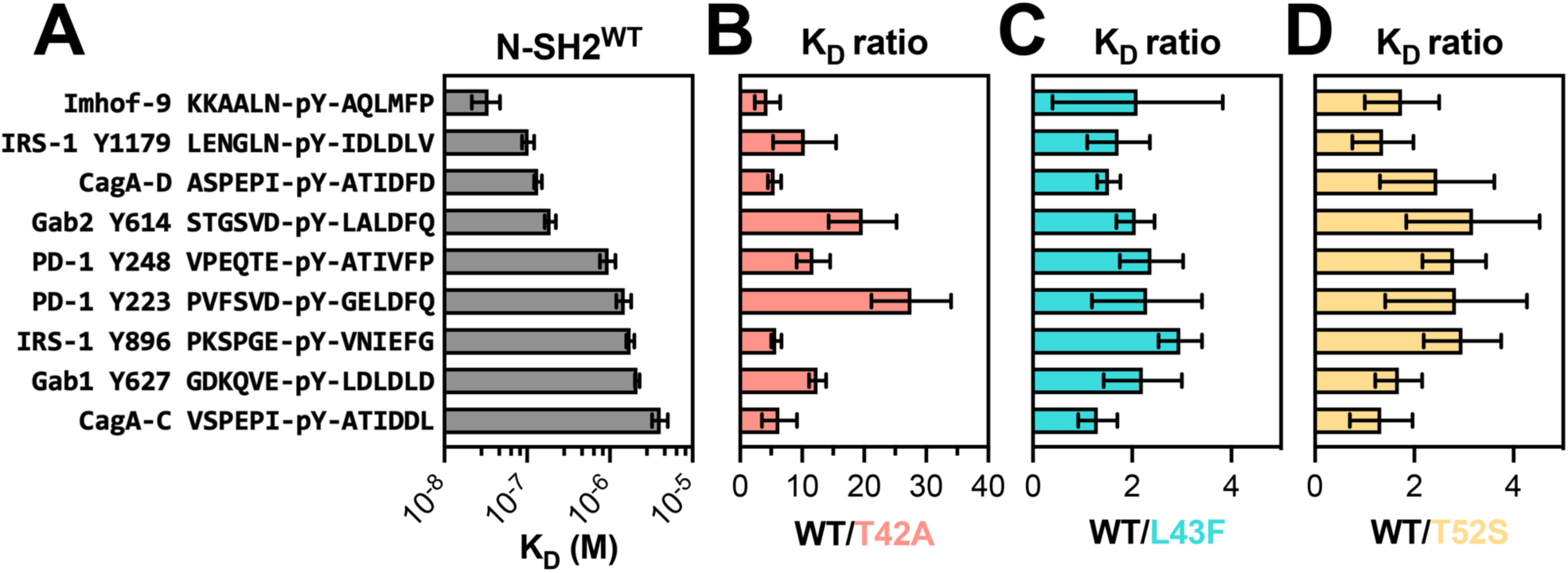
K_D_ measurements reveal sequence-specific enhancement of binding affinity in N-SH2^T42A^. (**A**) Measured binding affinities of N-SH2^WT^ against peptides derived from various known SHP2 interactors. (**B**) Fold-change in K_D_ for N-SH2^T42A^ compared to N-SH2^WT^, for each of the peptides shown in panel (A). (**C**) Same as (B), but for N-SH2^L43F^. (**D**) Same as (B), but for N-SH2^T52S^. For (**A**)-(**D**), N = 3-4 independent protein, peptide, and fluorescent peptide titrations. Source data and p-values can be found in Table S2.

For C-SH2 domain mutants R138Q and E139D, we first measured binding against two fluorescent phosphopeptides: one derived from a known binding site on PD-1 (pTyr^248^), as well as the designed ligand Imhof-9 (**Figure S2D and Table S2**) (32, 33). As expected, C-SH2^R138Q^ binding to phosphopeptides was severely attenuated (**Figure S2D**), and this mutant was therefore excluded from further binding analyses. C-SH2^E139D^ binding was comparable to C-SH2^WT^ binding against the two fluorescent phosphopeptides (**Figure S2D**) and against the 9 unlabeled peptides used for N-SH2 binding assays (**Figure S2E and Table S2**). Collectively, these N-SH2 and C-SH2 binding experiments with a small panel of peptides suggest that N-SH2^T42A^ is unique among the SH2 mutants in its impact on both phosphopeptide binding affinity and specificity.

### Human phosphopeptide profiling reveals the scope of specificity differences in SHP2 SH2 domain mutants

Next, we characterized the sequence specificity of the SH2 mutants relative to their wild-type counterparts in a large-scale, unbiased screen. This method entails bacterial display of a peptide library, enzymatic phosphorylation of the displayed peptides, selective enrichment using SH2-coated beads, and deep sequencing of the peptide-coding DNA (34). For each peptide in a library, an enrichment score is calculated from the deep sequencing data as the frequency of the peptide in the SH2-enriched sample divided by the frequency of the peptide in the unenriched input sample (34). For this study, we used two largely non-overlapping libraries, both encoding known human phosphosites. The pTyr-Var Library contains 3,065 sequences corresponding to wild-type tyrosine phosphosites, with an additional 6,833 sequences encoding disease-associated point mutations, natural polymorphisms, or control mutations (34). The Human pTyr Library consists of 1,916 sequences derived from known phosphorylation sites in the human proteome, along with another 617 control mutants (35).

We profiled N-SH2^WT^, N-SH2^T42A^, N-SH2^L43F^, N-SH2^T52S^, C-SH2^WT^, and C-SH2^E139D^, against both libraries described above. The libraries were screened separately, but under identical conditions, and the spread of peptide enrichment scores was similar across both libraries. Thus, the results of both screens were combined for the analyses described below. We omitted sequences from our analysis that contained more than one tyrosine residue, yielding 9,281 relevant sequences across both libraries. For most phosphopeptides the screens showed a strong correlation between enrichment scores for the wild-type SH2 domain and the corresponding SH2 mutants. However, some phosphopeptides had larger enrichment scores for the mutant N-SH2 domains when compared to N-SH2^WT^ (**Figure 3A-C and Table S3**). This effect was strongest for N-SH2^T42A^, both in magnitude and in number of phosphopeptides that were disproportionately enriched in the N-SH2^T42A^ screens. In the C-SH2 domain screens, C-SH2^E139D^ showed slightly weakened binding to some peptides when compared to C-SH2^WT^ (**Figure 3D and Table S3**), in contrast to our binding affinity measurements (**Figure S2D,E**). This result is in alignment with previous work showing a change in binding preferences for C-SH2^E139D^, and it reinforces the importance of screening a large number of peptides for an unbiased assessment of specificity (14).

**Figure 3.**
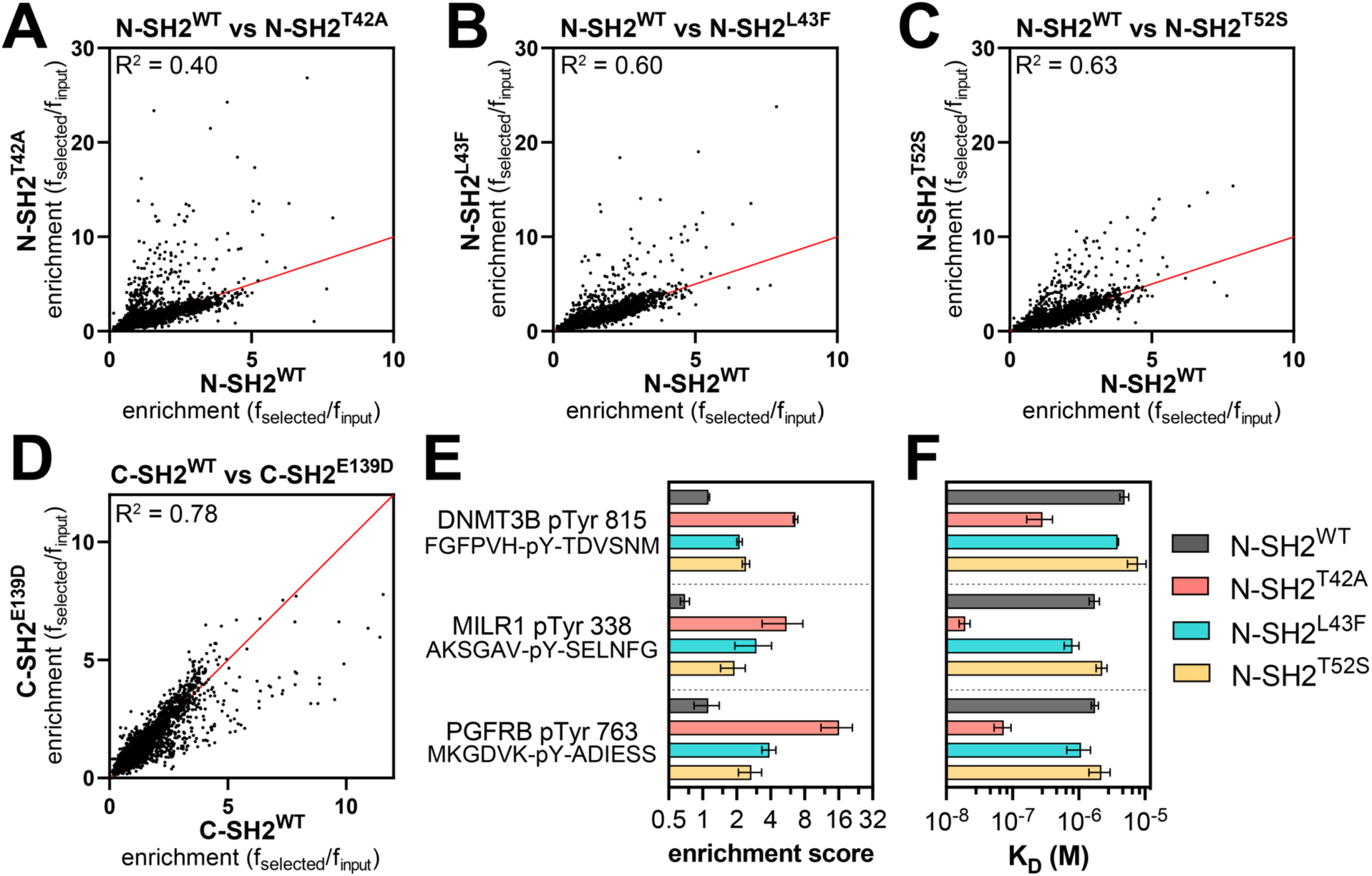
Peptide library screens identify sequences with enhanced N-SH2^T42A^ binding. (**A-C**) Comparison of enrichment scores for N-SH2^WT^ and each N-SH2 mutant. (**D**) Comparison of enrichment scores for C-SH2^WT^ and C-SH2^E139D^. The red line denotes x = y. pTyr-Var Library screens (N = 2), Human pTyr Library screens (N = 3). Source data for panels (A)-(D) can be found in Table S3. (**E**) Enrichment scores from peptide display screens for three representative peptides that showed enhanced binding to N-SH2^T42A^ relative to N-SH2^WT^. (**F**) Binding affinity measurements for the three peptides shown in panel (E) (N = 3).

To validate the stark difference between N-SH2^WT^ and N-SH2^T42A^ in our screens, we selected three sequences for fluorescence polarization binding affinity measurements (**Figure 3E**). One of these peptides, DNMT3B pTyr^815^, was derived from DNA methyltransferase 3β, a protein that is not known to interact with SHP2. The second peptide, PGFRB pTyr^763^, was derived from the platelet-derived growth factor receptor β, which is known to interact with SHP2 through this phosphosite (36). The third peptide, MILR1 pTyr^338^, was derived from a known SHP2 binding site on the mast cell immunoreceptor Allegrin-1 (37). Competition fluorescence polarization assays with these peptides revealed large differences in binding affinity between N-SH2^WT^ and N-SH2^T42A^, as predicted by the screens (**Figure 3F and Table S2**). N-SH2^T42A^ bound 17-fold tighter to DNMT3B pTyr^815^ than N-SH2^WT^ and 24-fold tighter to PGFRB pTyr^763^. The difference in binding affinity between N-SH2^WT^ and N-SH2^T42A^ was largest for MILR1 pTyr^338^, for which the mutation caused a 90-fold enhancement.

### SHP2 N-SH2^WT^ and N-SH2^T42A^ display distinct position-specific sequence preferences

Next, we examined the sequence features of the peptides enriched in the screens with each N-SH2 domain. Our peptide libraries collectively contain 392 sequences lacking tyrosine residues, which serve as negative controls in our screens. Less than 2% of the negative control peptides had enrichment scores above of 3.2, and so we used this value as a stringent cutoff to identify true binders in each screen, as done previously (34). We identified 168 enriched sequences for N-SH2^WT^ and approximately 250 enriched sequences for each of the N-SH2 mutants, indicative of overall tighter binding by the mutants (**Figure 4A, Figure S3A, and Table S3**). Consistent with its unique change in binding specificity, the enriched peptide set for N-SH2^T42A^ had less overlap with that of N-SH2^WT^ when compared with N-SH2^L43F^ or N-SH2^T52S^. Probability sequence logos, derived by comparing the amino acid composition of these enriched peptide sets to the full library, showed that N-SH2^T42A^ had the most distinctive sequence preferences of all four N-SH2 variants (**Figure S3B-E**) (38).

**Figure 4.**
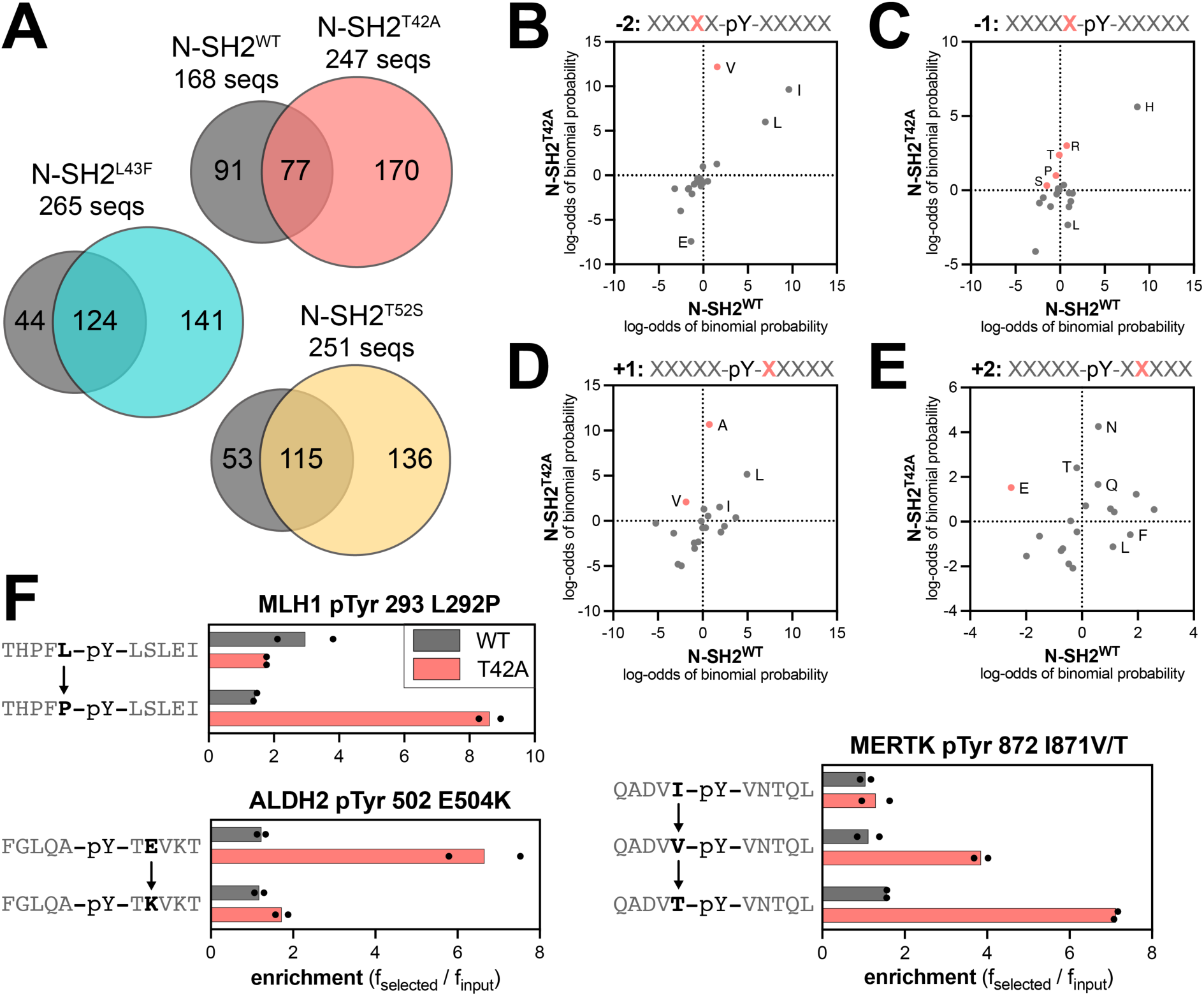
Analysis of enriched sequence features reveals specificity differences at several positions. (**A**) Overlap in sequences enriched by each N-SH2 domain above an enrichment score cutoff of 3.2. (**B**) Log-transformed probabilities of amino acid enrichment at the -2 position relative to the pTyr residue, derived from peptide display screens with N-SH2^WT^ and N-SH2^T42A^. (**C**) Same as panel (B), except at the -1 position. (**D**) Same as panel (B), except at the +1 position. (**E**) Same as panel (B), except at the +2 position. (**F**) Enrichment scores for representative sets of peptides from the pTyr-Var Library screens that highlight different sequence preferences for N-SH2^WT^ and N-SH2^T42A^. N = 2 independent library transformations and screens.

Due to the small number of enriched sequences in the N-SH2^WT^ screens, the corresponding sequence logo has low signal-to-noise ratio. Even so, the logo highlights several hallmarks of the SHP2 N-SH2 domain, such as a preference for a -2 Ile, Leu or Val, -1 His, +3 hydrophobic residue, and +5 Phe or His (19, 32). As expected, the N-SH2 mutants share many of these features with N-SH2^WT^ (**Figure S3B-E**). However, we observed distinct changes in specificity at positions closest to the pTyr residue. N-SH2^T42A^ prefers the smaller Val over Ile and Leu on the -2 position (**Figure 4B and Figure S3B,C**). At the -1 position, although His is strongly favored for all four SH2 domains, N-SH2^T42A^ had broadened tolerance of other amino acids, including Pro and polar residues Gln, Ser, Thr, and Arg (**Figure 4C and Figure S3B,C**). At the +1 position, N-SH2^WT^ favors large hydrophobic residues (Leu, Ile, or Phe), as well as His and Asn. By contrast, Ala is the dominant preference for N-SH2^T42A^ and is also enriched for N-SH2^L43F^ and N-SH2^T52S^, but to a lesser extent (**Figure 4D and Figure S3B-E**). At the +2 position, we found that N-SH2^T42A^ has a unique enhanced preference for hydrophilic residues. One notable difference, discussed in subsequent sections, is a switch for +2 Glu from a disfavored residue for N-SH2^WT^ to a favored residue for N-SH2^T42A^ (**Figure 4E and Figure S3B,C**). Finally, at the +3 residue, each N-SH2 variant shows a slightly different preference for specific hydrophobic residues, with N-SH2^T42A^ strongly preferring Leu (**Figure S3B-E**).

The sequence logos represent the position-specific amino acid preferences of each N-SH2 variant, without taking into account the surrounding context. The pTyr-Var Library used in the screens encodes wild-type and point-mutant sequences derived from human phosphorylation sites, providing an internal control for sequence-specific mutational effects. Upon closer inspection of individual hits for N-SH2^WT^ and N-SH2^T42A^, we identified several sets of sequences that corroborate the overall preferences described above. These include an enhanced preference in N-SH2^T42A^ for Pro or Thr at the -1 position over Leu or Ile and a strong preference for a +2 Glu residue (**Figure 4F**).

To more comprehensively analyze sequence preferences in a physiologically-relevant sequence context, we generated a saturation mutagenesis library based on the sequence surrounding PD-1 pTyr^223^, and we screened this library against the N-SH2^WT^ and N-SH2^T42A^ using bacterial peptide display. The immunoreceptor tyrosine-based inhibitory motif (ITIM) surrounding PD-1 pTyr^223^ was chosen because this was a sequence for which we observed a large change in binding affinity between N-SH2^WT^ and N-SH2^T42A^ (**Figure 2B**). Due to the relatively weak binding of N-SH2^WT^ to the wild-type PD-1 pTyr^223^ peptide, the differentiation of neutral mutations and loss-of-function mutations was poor in our screen (**Figure S3F-G and Table S4**). However, we could confidently detect gain-of-function mutations for N-SH2^WT^. For N-SH2^T42A^, the overall tighter binding affinity allowed for reliable measurement of both gain- and loss-of-function point mutations on this peptide (**Figure S3F-G and Table S4**).

Our results show that the two domains have modestly correlated binding preferences with respect to this scanning mutagenesis library (**Figure S3G**). The -1 Asp and +1 Gly residues in the ITIM sequence are suboptimal for both N-SH2^WT^ and N-SH2^T42A^, as most substitutions at these positions enhance binding. However, differences were observed in which mutations were tolerated by each SH2 domain at these positions (**Figure S3F**). For example, the substitution of the +1 Gly to Ala or Thr is favored by N-SH2^WT^, consistent with previous studies (19), but large hydrophobic residues are also favorable for N-SH2^WT^ at this position. By contrast, N-SH2^T42A^ strongly disfavors a +1 Trp and Phe. This recapitulates our analysis of sequences enriched in the human phosphopeptide library screens, where we observed a N-SH2^WT^ preference for larger residues (Leu, Ile, Phe), whereas N-SH2^T42A^ had a strong preference for the smaller alanine (**Figure 4D**). Also consistent with our analysis of the human phosphopeptide screens, most substitutions at the -2 Val or +2 Glu in the ITIM are heavily disfavored by N-SH2^T42A^ (**Figure S3F**). Taken together, our experiments with the human phosphosite libraries and the scanning mutagenesis library highlight consistent differences in the sequence preferences of N-SH2^WT^ and N-SH2^T42A^, suggestive of distinct modes of phosphopeptide engagement by these two domains.

### The T42A mutation enhances binding by remodeling the N-SH2 phosphotyrosine binding pocket

Several structural explanations for tighter phosphopeptide binding by N-SH2^T42A^ have been postulated previously (14, 17, 39–41). Crystal structures of N-SH2^WT^ bound to different peptides show that the hydroxyl group of the Thr^42^ side chain hydrogen bonds to a non-bridging oxygen atom on the phosphotyrosine moiety of the ligand (**Figure 5A**) (33, 42–45). The loss of this hydrogen bond in the T42A mutant is thought to be counterbalanced by enhanced hydrophobic interactions between the pTyr phenyl ring and Ala^42^ (14, 17, 39–41), but this cannot explain differences in the recognition of the surrounding peptide sequence, which is over 10 Å away. Many SH2 domains have a hydrophobic residue (Ala, Val, or Leu) at the position corresponding to Thr^42^ (**Figure S4A**) (46), but the impact of this residue on sequence specificity has not been systematically explored. Here, we used molecular dynamics (MD) simulations to examine how the T42A mutation impacts SHP2 N-SH2 peptide engagement (39, 47, 48).

**Figure 5.**
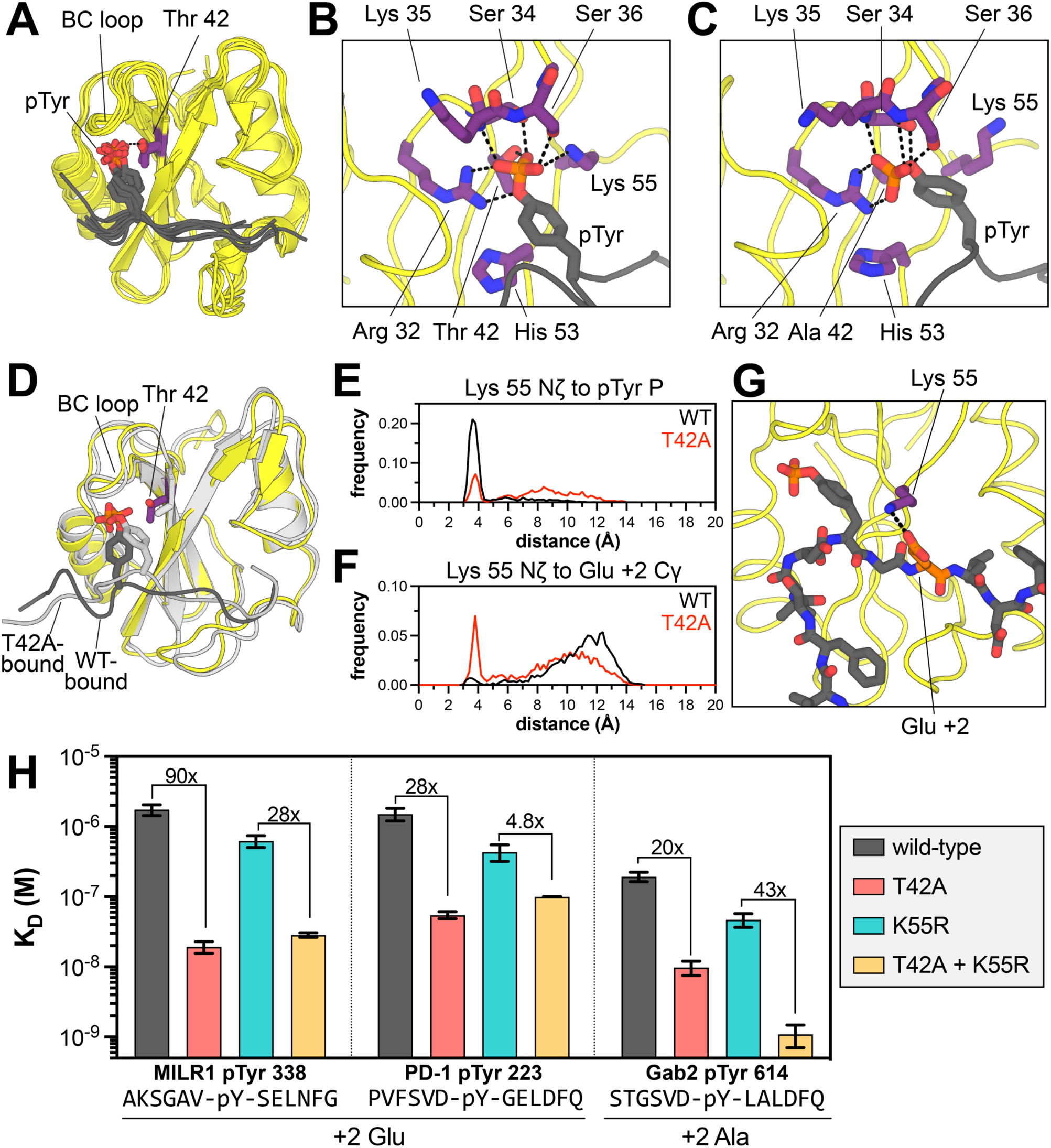
Structural impact of the T42A mutation on phosphotyrosine and proximal sequence recognition. (**A**) Hydrogen bonding of Thr^42^ in SHP2 N-SH2^WT^ to the phosphoryl group of phosphopeptide ligands in several crystal structures (PDB: 6ROY, 1AYA, 1AYB, 3TL0, 5DF6, 5X94, and 5X7B). (**B**) Representative structures of (**B**) N-SH2^WT^ and (**C**) N-SH2^T42A^ bound to the PD-1 pTyr^223^ (ITIM) peptide at the ends of respective trajectories. (**D**) Overlay of the states shown in panels B and C. The N-SH2^WT^ state is in yellow with a dark-gray ligand. The N-SH2^T42A^ state is in light gray, with a light gray ligand. (**E**) Distribution of distances between the Lys^55^ Nζ atom and the phosphotyrosine phosphorus atοm in simulations of the PD-1 pTyr^223^ peptide bound to N-SH2^WT^ (black) or N-SH2^T42A^ (red). (**F**) Distribution of distances between the Lys^55^ Nζ atom and the +2 Glu Cδ atom in simulations of the PD-1 pTyr^223^ peptide bound to N-SH2^WT^ (black) or N-SH2^T42A^ (red). (**G**) An ion pair between Lys^55^ and the +2 Glu residue (Glu 225) in the PD-1 pTyr^223^ (ITIM) peptide, frequently observed in N-SH2^T42A^ simulations. (**H**) Peptide-specific effects of the T42A mutation in the presence and absence of the K55R mutation. N = 3-5 independent titrations. All fold-changes and respective p-values can be found in Table S2.

We carried out simulations of SHP2 N-SH2^WT^ and N-SH2^T42A^ in the apo state and bound to five different phosphopeptide ligands (PD-1 pTyr^223^, MILR1 pTyr^338^, Gab2 pTyr^614^, IRS-1 pTyr^896^, and Imhof-9). Each system was simulated three times, for 1 μs per run. We first calculated the per-residue root mean squared fluctuation (RMSF) in each system (**Figure S5A**). Simulations of the peptide-bound state showed rigidification of the BC loop (residues 32-40) relative to apo-state simulations. This loop is responsible for coordinating the phosphoryl moiety through a series of hydrogen bonds. Closer inspection of BC loop interactions in the N-SH2^T42A^ simulations revealed substantial reorganization of the hydrogen bond network around the phosphoryl group, which alters the positioning of phosphotyrosine within the N-SH2 binding pocket. In every N-SH2^WT^ simulation, Thr^42^ made a persistent hydrogen bond with a non-bridging oxygen on the phosphoryl group, which constrains the orientation of the phosphotyrosine residue (**Figure 5B and Figure S5B,C**). By contrast, in almost every N-SH2^T42A^ simulation, where the phosphoryl group was not tethered to Thr^42^, the phosphotyrosine residue relaxed into a new orientation characterized by a distinct hydrogen bond network (**Figure 5C**, *additional structural details in* **Figure S5B-H**).

### T42A-dependent changes in phosphotyrosine engagement drive changes in sequence recognition

A major consequence of the reshuffling of hydrogen bonds between N-SH2^WT^ and N-SH2^T42A^ is that the phosphotyrosine residue was positioned slightly deeper into the ligand binding pocket of the mutant SH2 domain. The phenyl ring moved distinctly closer to residue 42 in the mutant simulations, presumably engaging in stabilizing hydrophobic interactions (**Figure 5D and Figure S5F**) (47, 48). Overall, the peptide main chain residues near the phosphotyrosine appeared closer to the body of the SH2 domain (**Figure 5D**). This likely alters side chain packing at the interface and may explain why N-SH2^T42A^ prefers smaller residues at the -2 position (Val over Leu/Ile) and +1 position (Ala over Leu/Ile) (**Figure 4**). Consistent with this, the Cα-to-Cα distance between the peptide +1 residue and Ile^54^, which lines the peptide binding pocket, was frequently shorter in N-SH2^T42A^ simulations than N-SH2^WT^ simulations (**Figure S5H**).

One striking difference between the N-SH2^WT^ and N-SH2^T42A^ simulations in the peptide-bound state was the positioning and movement of Lys^55^. In crystal structures and in our N-SH2^WT^ simulations, the Lys^55^ ammonium group interacted with the phosphotyrosine phosphoryl group or engaged the phenyl ring in a cation-π interaction (**Figure 5B,E and Figure S6A**) (33, 42–45). In the N-SH2^T42A^ simulations, the phosphoryl group rotated away from Lys^55^ and engaged the BC loop and Arg^32^, thereby liberating the Lys^55^ side chain (**Figure 5C,E and Figure S6A**). This shift altered the electrostatic surface potential of N-SH2^T42A^ in the peptide binding region when compared to N-SH2^WT^ (**Figure S6B**). In some simulations, the Lys^55^ side chain ion paired with the Asp^40^ side chain (**Figure S6C**). For the N-SH2^T42A^ simulations with the PD-1 pTyr^223^ peptide, we observed substantial sampling of a distinctive state, where the Lys^55^ side chain formed an ion pair with the +2 Glu residue (PD-1 Glu^225^) (**Figure 5F,G**). This interaction was not observed in the N-SH2^WT^ simulations, and indeed, our peptide display screens showed enhanced preference for a +2 Glu by N-SH2^T42A^ over N-SH2^WT^ (**Figure 4E-G**). Other peptides had Asp and Glu residues at nearby positions, but stable ion pairs between these residues and Lys^55^ were not observed in our simulations.

Only three human SH2 domains have an Ala and Lys at the positions that are homologous to residues 42 and 55 in SHP2 N-SH2: the SH2 domains of Vav1, Vav2, and Vav3 (**Figure S4B,C**) (46). Experimental structures of the Vav2 SH2 domain bound to phosphopeptides show that the Lys^55^-analogous lysine residue can form electrostatic interactions with acidic residues at various positions on the peptide ligands, including the +2 residue (**Figure S6D**) (49, 50). Furthermore, Vav-family SH2 domains are known to prefer a +2 Glu on their ligands (51), further corroborating a role for SHP2 Lys^55^ in substrate selectivity in an T42A-context.

We hypothesized that Lys^55^ plays an important role in the specificity switch caused by the T42A mutation. Thus, we conducted a double-mutant cycle analysis in which we examined the effect of the T42A mutation in the presence and absence of a K55R mutation, measuring binding affinities to phosphopeptides with and without a +2 Glu (**Figure 5H**). Many SH2 domains have an arginine at position 55 (**Figure S4B**). This Arg residue forms a cation-π interaction with the pTyr phenyl ring, interacts with the phosphoryl group, or engages a conserved acidic residue at the end of the BC loop. All of these interactions with arginine would likely be tighter than analogous interactions with Lys^55^ and may persist in a T42A context. Indeed, the K55R mutation, on its own, enhanced binding to all measured phosphopeptides, with an effect ranging from 2-fold to 7-fold (**Figure 5H, Figure S6E** *gray vs. cyan,* **and Table S2**). As discussed earlier, the T42A mutation on its own enhanced binding anywhere from 4-fold to 90-fold (**Figure 5H, Figure S6E** *gray vs. red*, and **Figure 2**).

For peptides with a +2 Glu, the effect of the T42A mutation was large in isolation, but was substantially diminished in presence of a K55R mutation (**Figure 5H, Figure S6F, and Table S2**). For example, for the MILR1 pTyr 338 peptide, the 90-fold enhancement of binding affinity caused by the T42A mutation was attenuated to 28-fold in a K55R background. For the PD-1 pTyr 223 peptide, which also has a +2 Glu, the T42A mutation alone enhanced binding 28-fold, but the effect of this mutation dropped to 4.3-fold for in the presence of K55R. The Gab2 pTyr 614 peptide has a similar sequence to the PD-1 pTyr 223 peptide, but with an Ala at the +2 position. For this peptide, the T42A mutation enhanced binding 20-fold, but in contrast to the PD-1 and MILR1 peptides, this effect was actually enhanced to 43-fold in the presence of the K55R mutation (**Figure 5H, Figure S6F, and Table S2**). For peptides lacking a +2 Glu that showed a small enhancement in binding affinity upon the T42A mutation, such as IRS-1 pTyr 1179 and Imhof-9, the T42A effect was not impacted by the K55R mutation (**Figure S6E**).

These experiments strongly suggest that Thr^42^ and Lys^55^ are energetically coupled (**Figure S6F**), and that the dramatic enhancement in binding affinity of N-SH2^T42A^ for peptides with a +2 Glu is, at least in part, dependent on the presence of Lys^55^. The sequence similarity between the PD-1 pTyr 223 and Gab2 pTyr 614 peptides suggests that other T42A-specific sequence preferences, independent of Lys^55^ and a +2 Glu, can also contribute to enhanced binding. These include the T42A preference for a -2 Val described above (**Figure 4B**). Overall, the simulations and experiments in these past two sections provide a structural explanation for how the T42A mutation remodels the ligand binding pocket of the N-SH2 domain, resulting in a change in peptide selectivity.

### T42A-dependent changes in N-SH2 specificity drive changes in SHP2 activation

Given that N-SH2 engagement is thought to be the main driver of SHP2 activation (10, 39), we hypothesized that the binding specificity changes caused by the T42A mutation would sensitize SHP2 to some activating peptides but not others. To assess enzyme activation, we measured the catalytic activity full-length SHP2^WT^ against the fluorogenic substrate DiFMUP, in the presence of the phosphopeptides used in our binding affinity measurements (**Figure 6A**). SHP2^WT^ activity was enhanced with increasing phosphopeptide concentration, demonstrating ligand-dependent activation (**Figure 6B**). The concentration of phosphopeptide required for half-maximal activation (EC_50_) was different for each phosphopeptide and correlated well with the binding affinity of the phosphopeptide for the N-SH2 domain (**Figure 6C and Table S5**), substantiating the importance of N-SH2 domain engagement for activation.

**Figure 6.**
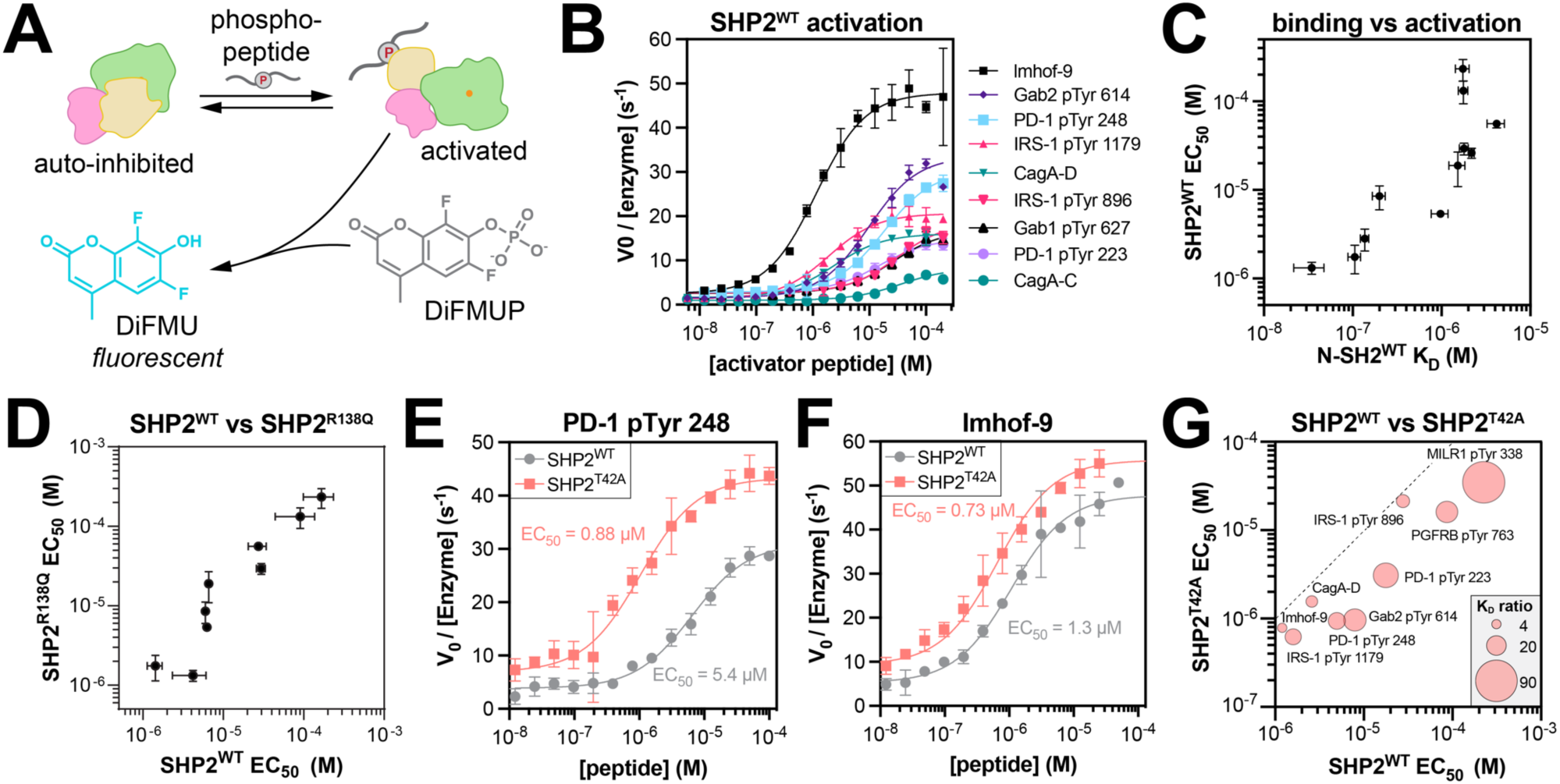
T42A-dependent changes in the activation of full-length SHP2. (**A**) Measurement of SHP2 activation by phosphopeptides. (**B**) Representative activation curves for SHP2^WT^ (N = 3-17). (**C**) Correlation between the EC_50_ of SHP2^WT^ activation by phosphopeptides and the K_D_ of those phosphopeptides for the N-SH2^WT^ domain (N = 3-4 for K_D_ values, N = 3-17 for EC_50_ values). (**D**) Activation EC_50_ values for SHP2^WT^ versus SHP2^R138Q^ (N = 3-17 for SHP2^WT^, N = 3-5 for SHP2^R138Q^). (**E**) Comparison of SHP2^WT^ and SHP2^T42A^ activation by PD-1 pTyr 248 (N = 3-4). (**F**) Comparison of SHP2^WT^ and SHP2^T42A^ activation by Imhof-9 (N = 6-17). (**G**) Bubble plot juxtaposing the EC_50_ values for activation of SHP2^WT^ and SHP2^T42A^ by nine peptides, alongside the fold-change in K_D_ for binding of those peptides to N-SH2^WT^ vs N-SH2^T42A^. The dotted line indicates where EC_50_ for SHP2^WT^ equals EC_50_ for SHP2^T42A^. Peptides with a large fold-change in binding affinity (larger bubble) have a large fold-change in EC_50_ values for SHP2^T42A^ over SHP2^WT^ (distance from dotted line). All EC_50_ values and p-values can be found in Table S5.

We also measured EC_50_ values for activation of full-length SHP2^R138Q^, which has negligible C-SH2 binding to phosphopeptides (**Figure S2C**). The EC_50_ values for SHP2^WT^ and SHP2^R138Q^ activation were strongly correlated, further supporting the notion that phosphopeptide binding to the N-SH2 domain, not the C-SH2 domain, is a major driver of SHP2 activation in our experiments (**Figure 6D and Table S5**). We note that some SHP2-binding proteins are bis-phosphorylated, unlike the mono-phosphorylated peptides tested in this work, and in that context, the C-SH2 domain can play an important role in activating SHP2 by localizing the N-SH2 domain to a binding site for which the N-SH2 otherwise has a weak affinity (52).

Previous reports showed that SHP2^T42A^ had enhanced activity when compared to wild-type SHP2 under saturating phosphopeptide concentration (14, 17). However, these experiments did not compare activation across multiple peptides. Thus, we measured activation of SHP2^T42A^ using our panel of phosphopeptides, allowing us to ascertain peptide-specific effects on the activation of SHP2^T42A^. For some peptides, the T42A mutation strongly shifted the EC_50_ to lower concentrations, whereas other activation curves were only marginally affected by the mutation (**Figure 6E,F and Table S5**). The peptides that showed a large enhancement in binding affinity to N-SH2^T42A^ over N-SH2^WT^ (**Figure 6G**, *large bubbles*) also showed the largest enhancement in activation (**Figure 6G**, *distance from dotted line*). These results demonstrate, for the first time, that the T42A mutation can sensitize SHP2 to specific activating ligands over others by altering N-SH2 binding affinity and specificity.

### The T42A mutation in SHP2 impacts its cellular interactions and signaling

All of the previous experiments were done using purified proteins and short phosphopeptide ligands. We next sought to determine if the effects of the T42A mutation could be recapitulated in a cellular environment with full-length proteins. First, we assessed the impact of the T42A mutation on phosphoprotein binding through co-immunoprecipitation experiments. We expressed myc-tagged SHP2^WT^ or SHP2^T42A^ in human embryonic kidney (HEK) 293 cells, along with a SHP2-interacting protein of interest and a constitutively active variant of the tyrosine kinase c-Src, to ensure phosphorylation of our interacting protein of interest. We chose Gab1, Gab2, and PD-1 as our proteins of interest, as these proteins play important roles in SHP2-relevant signaling pathways (28, 33, 53–56). For all three proteins, we found that SHP2^T42A^ co-immunoprecipitated more with the interacting protein than SHP2^WT^ (**Figure 7A and Figure S7A,B**). Anti-phosphotyrosine staining confirmed that these immunoprecipitated proteins were phosphorylated. We also examined whether the enhanced co-immunoprecipitation observed for SHP2^T42A^ was indeed the result of increased intrinsic N-SH2 binding affinity, as opposed to an increased occupancy of the open conformation of the full-length protein, which exposes the SH2 domains. To test this, we compared SHP2^E76K^, which predominantly occupies the open confirmation, to the SHP2^T42A+E76K^ double mutant. More Gab2 was co-immunoprecipitated with SHP2^T42A+E76K^ than with SHP2^E76K^ (**Figure 7A**), suggesting that T42A and E76K enhance binding through non-redundant mechanisms.

**Figure 7.**
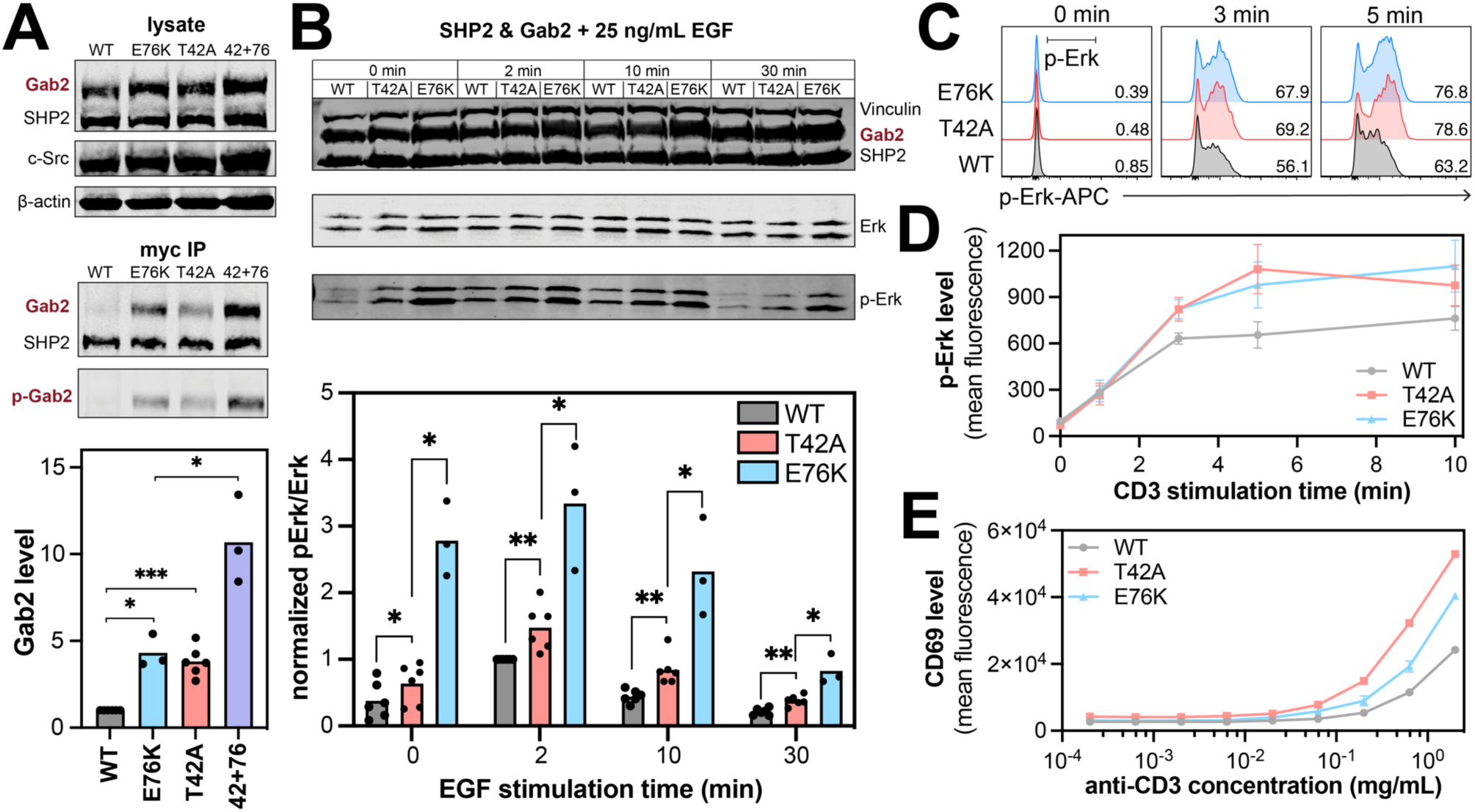
Enhanced protein-protein interactions and downstream signaling by SHP2^T42A^. (**A**) Gab2 co-immunoprecipitation with SHP^WT^, SHP2^E76K^, SHP2^T42A^ and SHP2^T42A+E76K^ (42+76). Relative Gab2 co-immunoprecipitation levels are normalized for expression and shown in the bar graph. (**B**) Phospho-Erk (p-Erk) levels in response to EGF stimulation in cells expressing Gab2 with either SHP2^WT^, SHP2^T42A^ SHP2^E76K^ (N = 3-6). The bar graphs below the blots show p-Erk levels, normalized to total Erk levels, relative to the highest p-Erk signal in the SHP2^WT^ time course (2 minutes). For all bar graphs, a paired, one-tailed t-test was used to test for significance (* denotes p < 0.05, ** denotes p < 0.01, *** denotes p < 0.001). (**C**) Representative histograms for Erk phosphorylation responses in wild-type, T42A, or E76K SHP2-expressing Jurkat variants stimulated with anti-CD3 antibody for the indicated time. The black bar in the first panel depicts the gate used to define the pErk+ population. (**D**) Geometric mean fluorescence intensity for phosho-Erk levels in SHP2^WT^, SHP2^T42A^ or SHP2^E76K^-expressing SHP2-deficient Jurkat variants (N = 3 stimulation replicates). (**E**) Geometric mean fluorescence intensity for CD69 levels in SHP2^WT^, SHP2^T42A^ or SHP2^E76K^-expressing SHP2-deficient Jurkat variants (N = 2 stimulation replicates). The data in panels **D** and **E** are representative of 5-6 biological replicates.

Next, we assessed whether enhanced SHP2 binding caused by the T42A mutation impacts downstream signaling. SHP2 is a positive regulator of Ras GTPases, and it can promote activation of the Ras/MAPK pathway downstream of receptor tyrosine kinases (57, 58). Its functions in this context can be mediated by the adaptor proteins Gab1 and Gab2 (59). Thus, we transfected SHP2^WT^ or SHP2^T42A^ into HEK 293 cells, along with Gab1 or Gab2. Cells were stimulated with epidermal growth factor (EGF), and Erk phosphorylation was analyzed as a marker of MAPK activation downstream of the EGF receptor. For the Gab2 experiment, we included SHP2^E76K^, which should have high levels of phospho-Erk, even in absence of stimulation. We observed a stimulation-time-dependent increase then decrease in phospho-Erk levels for all three mutants. Notably, the response was stronger and longer for SHP2^T42A^ than for SHP2^WT^ samples (**Figure 7B and Figure S7C,D**). A previous study using the isolated N-SH2^T42A^ and N-SH2^WT^ domains showed that the T42A mutation increased co-immunoprecipitation of growth factor signaling proteins from HeLa cell lysates, but effects on signaling were not directly assessed (60). Our results directly demonstrate that tighter binding to phosphoproteins by full-length SHP2^T42A^ can enhance downstream Ras/MAPK signaling.

Since SHP2 is a ubiquitously expressed protein, we investigated the functional effects of the T42A mutation in different cellular context, namely in T cells. We prepared SHP2 knock-out Jurkat T cells (**Figure S7E**), and reconstituted them with either SHP2^WT^, SHP2^T42A^ or SHP2^E76K^. The wild-type and mutant SHP2-expressing cells were labeled with different concentrations of the fluorescent dye CellTrace Violet and pooled (**Figure S7F**). Then, the cells were stimulated with a CD3-specific antibody to activate T cell receptor-CD3 complexes, and signal transduction was evaluated by measuring phospho-Erk levels using flow cytometry (**Figure 7C**). In agreement with our EGF stimulation experiments in HEK 293 cells, phospho-Erk levels in T cells were higher with SHP2^T42A^ and SHP2^E76K^ when compared with SHP2^WT^ (**Figure 7D and Figure S7G**). Next, we studied the effects of the T42A mutation on full T cell activation, as measured by expression of the activation marker CD69 (**Figure 7E and Figure S7H**). SHP2^T42A^ showed enhanced T cell activation at lower doses of CD3 stimulation. Interestingly, SHP2^E76K^ had an intermediate effect between SHP2^WT^ and SHP2^T42A^. SHP2 can participate in signaling pathways that both promote T cell activation (e.g. by activating the MAPK pathway) and suppress T cell activation (e.g. through the co-inhibitory receptor PD-1) (61, 62). Thus, we hypothesize that the reduced activation by SHP2^E76K^ when compared with SHP2^T42A^ may be a consequence of SHP2^E76K^ tapping into both positive and negative regulatory pathways, without the need for SH2 ligand binding. By contrast, SHP2^T42A^ may only be able to engage in negative regulatory pathways when an appropriate SH2 binding partner is presented.

## Discussion

The protein tyrosine phosphatase SHP2 is involved in a broad range of signaling pathways and has critical roles in development and cellular homeostasis. The most well-studied mutations in SHP2 disrupt auto-inhibitory interactions between the N-SH2 and PTP domain, thereby hyperactivating the enzyme and enhancing signaling. These mutations partly or fully decouple SHP2 activation from SHP2 localization – in this context, SHP2 no longer requires recruitment to phosphoproteins via its SH2 domains for full activation. By contrast, mutations outside of the N-SH2/PTP interdomain interface operate through alternative pathogenic mechanisms, and can have distinct outcomes on cellular signaling (11). In this study, we have characterized some of the mutations in this category, focusing on mutations in the phosphopeptide binding pockets of the SH2 domains. Our mutations of interest cause a wide range of disease phenotypes: Noonan Syndrome, juvenile myelomonocytic leukemia, acute lymphocytic leukemia, melanoma, and non-syndromic heart defects (5, 11, 15, 16). Most of these mutations do not have extensive biochemical data quantitatively characterizing their effects on phosphopeptide binding specificity or downstream cell signaling.

Here, we report a sequence-specific enhancement of binding affinity to tyrosine-phosphorylated ligands caused by the T42A mutation in the N-SH2 domain of SHP2. Previous studies have reported increased binding affinity by the T42A mutation, but the change in sequence specificity was not known, nor was the effect of biasing SHP2 activation toward certain ligands (14, 40, 60). Our insights into N-SH2^T42A^ specificity may reflect enhancements in measurement accuracy and library size for our SH2 specificity profiling platform relative to previous methods (34). Critically, our findings are supported by biochemical and biophysical experiments with a large panel of physiologically relevant peptides and proteins, whereas most studies analyzing SHP2 SH2 mutants have focused on a single peptide at a time. Of note, the appearance of +2 Glu and -2 Val preferences in our N-SH2^T42A^ specificity screens were present in a sequence logo generated from previous peptide microarray data (14), but as those findings were subtle, the N-SH2^T42A^ binding profile was reported as similar to N-SH2^WT^ in that study.

We have also demonstrated functional and cellular consequences of the T42A mutation in SHP2. Specifically, this mutation causes SHP2 to bind tighter to certain phosphoproteins and is more strongly activated as a result. We show that this enhanced binding and activation translates to downstream effects, such as increased intensity and duration of MAPK signaling in two different cell lines. This is in agreement with an affinity purification mass spectrometry study comparing SHP2 N-SH2^WT^ and N-SH2^T42A^ which found mutation-dependent changes in their interaction networks in mammalian cells (60). In that study, N-SH2^T42A^ showed increased interactions with growth factor signaling proteins, including those involved in MAPK signaling. Our comparisons of SHP2^T42A^ with SHP2^E76K^ suggest that these mutations alter SHP2 function through distinct mechanisms. What remains unknown is how the biased interaction specificity of the T42A mutant impacts SHP2 signaling when compared with other mutations that simply alter binding affinity but not specificity. Our findings suggest that SHP2^T42A^ will be hyperresponsive to certain upstream signals (e.g. phosphorylated Gab1 and Gab2), but not to others. Further studies will be needed to more broadly assess T42A-induced changes in cell signaling, in order to fully understand the pathogenic mechanism of this variant.

Much remains unknown about the other mutations explored in this study. To our knowledge, no biochemical or cell biological characterization has previously been reported for the L43F mutation (15). Here, we showed a mild increase in basal activity of full-length SHP2^L43F^ and a corresponding decrease in melting temperature, suggesting that this mutation slightly destabilizes the auto-inhibited state. L43F also causes a slight increase in binding affinity to phosphopeptides. The T52S mutation has previously been reported to change binding affinity to Gab2, consistent with our data (63). We did not observe any large changes in sequence specificity, with the exception of a preference for smaller Ala over bulkier residues at the +1 position. In our simulations with N-SH2^WT^ and N-SH2^T42A^, we observed that the methyl group of Thr^52^ interacts with the side chain on the +1 residue of the peptide, which could explain how T52S alters the +1 preference. The R138Q mutation severely disrupts phosphopeptide binding to the C-SH2 domain. The molecular basis for the pathogenic effects of the E139D mutation remains elusive. While many studies have addressed its binding affinity and specificity, their results are ambiguous (14, 17, 60). Our high-throughput screens indicate weakened binding of some peptides to C-SH2^E139D^ when compared to C-SH2^WT^. Consistent with previous work, our basal activity measurements with full-length SHP2 show that the E139D mutation is activating, suggesting an undefined regulatory role for the C-SH2 domain that may be unrelated to ligand binding (17, 22).

Most disease-associated mutations that alter the functions of cell signaling proteins do so by disrupting their intrinsic regulatory capabilities – for SHP2, pathogenic mutations cluster at the auto-inhibitory interface between the N-SH2 and PTP domains and hyperactivate the enzyme by disrupting interdomain interactions. There is increasing evidence that mutations can also rewire signaling pathways by changing protein-protein interaction specificity (64–66). This has been demonstrated most clearly for protein kinases, where mutations have been identified that alter substrate specificity (64). It is noteworthy that not all specificity-determining residues in kinases are located directly in the ligand/substrate binding pocket, raising the possibility that distal mutations may allosterically alter sequence specificity (67, 68). A similar paradigm has been suggested for SH2 domains, where distal mutations may rewire interactions, however the position corresponding to Thr^42^ in SHP2 has not been implicated as a determinant of specificity (67). The biochemical and structural analyses presented in this paper reveal an unexpected outcome of the pathogenic T42A mutation, where ligand selectivity is altered over 10 Å from the mutation site. Notably, mutations at the analogous Thr residue in other SH2 domains have also been implicated in disease, substantiating the significance of this residue (69–71). More broadly, our results illustrate the importance of considering the structural plasticity of signaling proteins when evaluating specificity and suggest that the functional consequences of many disease-associated mutations could be misclassified if evaluated solely based on their locations in static protein structures.

## Materials and Methods

Key resources, including cell lines, plasmids, oligonucleotide primers, peptides, and proteins, are listed in **Table S6**. Detailed materials and methods can be found in the supplementary information appendix. These include details of protein and peptide production, binding and activity measurements, bacterial peptide display screens, cell culture experiments, and molecular dynamics simulations.

## Acknowledgements

We would like to thank the members of the Shah lab for their scientific insights and helpful discussions. We thank Jeanine Amacher and Marko Jovanovic for their constructive feedback on this manuscript.

## Author contributions

AEV conceived, designed, performed, analyzed, and interpreted the experiments; performed statistical analysis; and wrote the manuscript. RV conceived, designed, performed, analyzed, and interpreted the experiments and wrote the manuscript. CAC, ZJ, PAH, and WLL performed and analyzed the experiments and edited the manuscript. DK performed and analyzed the MD simulations and edited the manuscript. NHS designed, analyzed, and interpreted the experiments, analyzed the MD simulations, and wrote the manuscript.

## Data and materials availability

All of the processed data, including catalytic efficiencies, binding affinities, EC_50_ values, and enrichment scores from the high-throughput specificity screens are provided as supplementary table files. The raw FASTQ sequencing files from specificity screens, source data from MD simulations, and processed MD trajectories are available as a Dryad repository (DOI: 10.5061/dryad.msbcc2g41). Plasmids and DNA libraries generated in this study will made freely available upon request. There are no restrictions to the availability of reagents generated in this study.

