## Supplementary Information and Figures for "The pathogenic T42A mutation in SHP2 rewires the interaction specificity of its N-terminal regulatory domain"

[†] These authors contributed equally.

#### ORCiDs

Anne E. van Vlimmeren: 0000-0003-0379-4945

Rashmi Voleti: 0000-0002-3705-7460

Cassandra A. Chartier: 0009-0004-2474-9937

Ziyuan Jiang: 0009-0003-6035-4126

Deepti Karandur: 0000-0002-6949-6337

Preston A. Humphries: 0000-0001-8736-6432

Wan-Lin Lo: 0000-0002-9074-6847

Neel H. Shah: 0000-0002-1186-0626

#### This SI Appendix contains:

Materials and Methods pg 2

Figure S1. Basal activity measurements and melting temperatures of various SHP2 mutants. pg 10

Figure S2. Binding affinities for SH2 domains. pg 11

Figure S3. Analysis of sequence-features for N-SH2 mutants. pg 12

Figure S4. Amino acid residues at key positions in human SH2 domains. pg 13

Figure S5. Measured structural parameters from MD simulations of SHP2 N-SH2$^{WT}$ or N-SH2$^{T42A}$. pg 14,15

Figure S6. The role of Lys$^{55}$ on T42A-dependent peptide recognition. pg 16

Figure S7. Cell-signaling consequences of the T42A mutation. pg 17

Supplementary References pg 18

### Materials and Methods

#### Key resources table

Key resources, including cell lines, plasmids, oligonucleotide primers, peptides, and proteins, are listed in **Table S6**.

#### Purification of SH2 domains

The SHP2 full-length, wild-type gene used as the template for all SHP2 constructs in this study was cloned from the pGEX-4TI SHP2 WT plasmid, which was a generous gift from Ben Neel (Addgene plasmid #8322) (1). SHP2 SH2 domains were cloned into a $His_6$-SUMO-SH2-Avi construct (2). C43(DE3) cells were transformed with plasmids encoding both the respective SH2 domain and the biotin ligase BirA. Cells were grown in LB supplemented with 50 µg/mL kanamycin and 100 µg/mL streptomycin at 37 °C until cells reached an optical density at 600 nm ($OD_{600}$) of 0.5. IPTG (1 mM) and biotin (250 µM) were added to induce protein expression and ensure biotinylation of SH2 domains, respectively. Protein expression was carried out at 18 °C overnight. Cells were centrifuged and subsequently resuspended in lysis buffer (50 mM Tris pH 7.5, 300 mM NaCl, 20 mM imidazole, 10% glycerol, and freshly added 2 mM β-mercaptoethanol). The cells were lysed using sonication (Fisherbrand Sonic Dismembrator), and spun down at 14,000 rpm for 45 minutes. The supernatant was applied to a 5 mL Ni-NTA column (Cytiva). The resin was washed with 10 column volumes lysis buffer and wash buffer (50 mM Tris pH 7.5, 50 mM NaCl, 20 mM imidazole, 10% glycerol, and freshly added 2 mM β-mercaptoethanol). The protein was eluted off the Ni-NTA column in elution buffer (50 mM Tris pH 7.5, 50 mM NaCl, 500 mM imidazole, 10% glycerol) and brought onto a 5mL HiTrap Q Anion exchange column (Cytiva). The column was washed using Anion A buffer (50 mM Tris pH 7.5, 50 mM NaCl, 1 mM TCEP). Protein elution off the column was induced through a salt gradient between Anion A buffer and Anion B buffer (50 mM Tris pH 7.5, 1 M NaCl, 1 mM TCEP). The eluted protein was cleaved at the $His_6$-SUMO tag by addition of 0.05mg/mL $His_6$-tagged Ulp1 protease at 4ºC overnight. This cleavage cocktail was flowed through a 2 mL Ni-NTA gravity column (ThermoFisher) to isolate the cleaved protein away from uncleaved protein and Ulp1. Finally, the cleaved protein was purified by size-exclusion chromatography on a Superdex 75 16/600 gel filtration column (Cytiva) equilibrated with SEC buffer (20 mM HEPES pH 7.4, 150 mM NaCl, and 10% glycerol). Pure fractions were pooled and concentrated, and flash frozen in liquid $N_2$ for long-term storage at −80 °C.

#### Purification of full-length SHP2 proteins

Full-length SHP2 variants were cloned into a pET28-His-TEV plasmid from the pGEX-4TI SHP2 WT plasmid (1). BL21(DE3) cells were transformed with the respective plasmids, and were grown in LB supplemented with 100 µg/mL kanamycin at 37 °C until cells reached an $OD_{600}$ of 0.5. IPTG (1 mM) was added to induce protein expression, which was carried out at 18 °C overnight. Cells were centrifuged and subsequently resuspended in lysis buffer (50 mM Tris pH 7.5, 300 mM NaCl, 20 mM imidazole, 10% glycerol, and freshly added 2 mM β-mercaptoethanol). The cells were lysed using sonication (Fisherbrand Sonic Dismembrator), and spun down at 14,000 rpm for 45 minutes. The supernatant was applied to a 5 mL Ni-NTA column (Cytiva). The resin was washed with 10 column volumes lysis buffer and wash buffer (50 mM Tris pH 7.5, 50 mM NaCl, 20 mM imidazole, 10% glycerol, and freshly added 2 mM β-mercaptoethanol). The protein was eluted off the Ni-NTA column in elution buffer (50 mM Tris pH 7.5, 50 mM NaCl, 500 mM imidazole, 10% glycerol) and brought onto a 5mL HiTrap Q Anion exchange column (Cytiva). The column was washed using Anion A buffer (50 mM Tris pH 7.5, 50 mM NaCl, 1 mM TCEP). Protein elution off the column was induced through a salt gradient between Anion A buffer and Anion B buffer (50 mM Tris pH 7.5, 1 M NaCl, 1 mM TCEP). The eluted protein was cleaved at the $His_6$-TEV tag by addition of 0.10 mg/mL of $His_6$-tagged TEV protease at 4 °C overnight. This cleavage cocktail was flowed through a 2 mL Ni-NTA gravity column (ThermoFisher) to separate the cleaved protein from uncleaved protein and TEV protease. Finally, the cleaved protein was purified by size-exclusion chromatography on a Superdex 200 16/600 gel filtration column (Cytiva) equilibrated with SEC buffer (20 mM HEPES pH 7.5, 150 mM NaCl, and 10% glycerol). Pure fractions were pooled and concentrated, and flash frozen in liquid $N_2$ for long-term storage at −80 °C.

### Synthesis and purification of peptides for in vitro activation and binding measurements

Several of the peptides used in this study were purchased from a commercial vendor (SynPeptide). The remaining peptides used for in vitro kinetic and binding assays were synthesized using 9-fluorenylmethoxycarbonyl (Fmoc) solid-phase peptide chemistry. All syntheses were carried out using the Liberty Blue automated microwave-assisted peptide synthesizer from CEM under nitrogen atmosphere, with standard manufacturer-recommended protocols. Peptides were synthesized on MBHA Rink amide resin solid support (0.1 mmol scale). Each Nα-Fmoc amino acid (6 eq, 0.2 M) was activated with diisopropylcarbodiimide (DIC, 1.0 M) and ethyl cyano(hydroxyamino)acetate (Oxyma Pure, 1.0 M) in dimethylformamide (DMF) prior to coupling. The coupling cycles for phosphotyrosine and the amino acid directly after it were done at 75 °C for 15 s, then 90 °C for 230 s. All other coupling cycles were done at 75 °C for 15 s, then 90 °C for 110 s. Deprotection of the Fmoc group was performed in 20% (v/v) piperidine in DMF (75 °C for 15 s then 90 °C for 50 s), except for the amino acid directly after the phosphotyrosine which had an additional initial deprotection (25 °C for 300 s). The resin was washed (4x) with DMF following Fmoc deprotection and after Nα-Fmoc amino acid coupling. All peptides were acetylated at their N-terminus with 10% (v/v) acetic anhydride in DMF and washed (4x) with DMF.

After peptide synthesis was completed, including N-terminal acetylation, the resin was washed (3x each) with dichloromethane (DCM) and methanol (MeOH), and dried under reduced pressure overnight. The peptides were cleaved and the side chain protecting groups were simultaneously deprotected in 95% (v/v) trifluoroacetic acid (TFA), 2.5% (v/v) triisopropylsilane (TIPS), and 2.5% water, in a ratio of 10 µL cleavage cocktail per mg of resin. The cleavage-resin mixture was incubated at room temperature for 90 minutes, with agitation. The cleaved peptides were precipitated in cold diethyl ether, washed in ether, pelleted, and dried under air. The peptides were redissolved in a 50% (v/v) water/acetonitrile solution and filtered from the resin.

The crude peptide mixture was purified using reverse-phase high performance liquid chromatography (RP-HPLC) on either a semi-preparatory C18 column (Agilent, ZORBAX 300SB-C18, 9.4 x 250 mm, 5 µm) with an Agilent HPLC system (1260 Infinity II), or a preparatory C18 column (XBridge Peptide BEH C18 Prep Column, 19 x 150 mm, 5 µm) with a Waters prep-HPLC system (Prep 150 LC System). Flow rate for purification was kept at 4 mL/min (semi-preparative) or 17mL/min (preparative) with solvents A (water, 0.1% (v/v) TFA) and B (acetonitrile, 0.1% (v/v) TFA). Peptides were generally purified over a 40 minute (semi-preparative) or 13 minute (preparative) linear gradient from solvent A to solvent B, with the specific gradient depending on the peptide sample. Peptide purity was assessed with an analytical column (Agilent, ZORBAX 300SB-C18, 4.6 x 150 mm, 5 µm) at a flow rate of 1 mL/min over a 0-70% B gradient in 30 minutes. All peptides were determined to be ≥95% pure by peak integration. The identities of the peptides were confirmed by mass spectroscopy (Waters Xevo G2-XS QTOF). Pure peptides were lyophilized and redissolved in 100 mM Tris, pH 8.0, as needed for experiments.

### Synthesis and purification of fluorescent peptides for in vitro binding measurements

The fluorescent peptides were prepared as described above for the unlabeled peptides, except for the coupling of aminohexanoic acid (AHX) and the fluorescein isothiocyanate (FITC) at the N-terminus, instead of an N-terminal acetyl. AHX (6 eq, 0.2 M) was activated with diisopropylcarbodiimide (DIC, 1.0 M) and ethyl cyano(hydroxyamino)acetate (Oxyma Pure, 1.0 M) in dimethylformamide (DMF) prior to coupling. The coupling cycle was done at 75 °C for 35 s then 90 °C for 575 s. Deprotection was performed as described previously.

After peptide synthesis was completed, including N-terminal AHX-labeling, the resin was washed (3x each) with dichloromethane (DCM) and methanol (MeOH) and dried under reduced pressure overnight. Then, a portion of the resin (0.025 mmol) was prepared for FITC labeling by swelling in DMF with agitation for 30 min. Excess DMF was removed and to the resin was added FITC (0.075 mmol, 3 eq.) and DIPEA (0.15mmol, 6 eq.) in DMF. This reaction was incubated at room temperature with agitation for 2 hours. After FITC labeling, the resin was washed (3x each) with dichloromethane (DCM) and methanol (MeOH) and dried under reduced pressure overnight. Finally, the peptides were cleaved and purified as previously described.

### Basal SHP2 catalytic activity measurements

Initial rate measurements for the SHP2-catalyzed dephosphorylation of 6,8-difluoro-4-methylumbelliferyl phosphate (DiFMUP) were conducted at 37 °C in DiFMUP buffer (60 mM HEPES pH 7.2, 150 mM NaCl, 1mM EDTA, 0.05% Tween-20). Reactions of 50 µL were set up in a black polystyrene flat bottom half area 96-well plate. A substrate concentration series of 31.25 µM, 62.5 µM, 125 µM, 250 µM, 500 µM, 1000 µM, 2000 µM and 4000 µM was used to determine $k_{cat}$ and $K_M$. Reactions were started by addition of appropriate amount of SHP2 wild-type and mutants (wild-type: 2.5 nM; T42A: 2.5 nM; L43F: 2.5 nM; T52S: 4 nM; E76K: 0.5 nM; R138Q: 3 nM, E139D: 2.5 nM). Emitted fluorescence at 455nm was recorded every 25 seconds in a span of 50 minutes using a BioTek Synergy Neo2 multi-mode reader.

Initial rate measurement for the SHP2-catalyzed dephosphorylation of *p*-nitrophenyl phosphate (pNPP) were conducted at 37 °C in pNPP buffer (10 mM HEPES, pH 7.5, 150 mM NaCl, 1 mM TCEP). Reactions of 75uL were set up in a black polystyrene flat bottom half area 96-well plate. A substrate concentration series of 12.8, 6.4, 3.2, 1.6, 0.8, 0.4, 0.2 and 0.1mM pNPP was used to determine $K_{cat}$ and $K_M$. Reactions were started by addition of appropriate amount of SHP2 wild-type and mutants (wild-type: 250 nM;T42A: 250 nM; L43F: 250 nM; T52S: 250 nM; E76K: 100 nM; R138Q: 250 nM; E139D: 250 nM).

In all cases, the linear region of the reaction progress curve was determined by visual inspection and fit to a line. These slopes were converted from absorbance or fluorescence units as a function of time to product formation as a function of time using standard curves measured with the reaction products (*p*-nitrophenol and 6,8-difluoro-7-hydroxy-4-methylcoumarin). Finally, these rates were corrected for enzyme concentration by dividing the values the concentration of enzyme used in the experiment to yield $V_0$ / [enzyme] in units of ($s^{-1}$). These corrected rates were plotted as a function of substrate concentration and fit to the Michaelis-Menten equation using non-linear regression to determine catalytic parameters. Experiments were generally repeated at least three times, and the average and standard deviation of all individual replicates are reported.

### Fluorescence polarization binding assays

SH2 domains were thawed in room temperature water and their absorbance at 280 nm was measured to determine concentration. SH2 domains were serially diluted 15 times in assay buffer (60 mM HEPES pH 7.2, 75 mM KCl, 75 mM NaCl,1 mM EDTA, 0.05% Tween-20), with a 2x starting concentrations generally in the low micromolar range. One well did not contain any SH2 domain. The fluorescent peptide was diluted to 2x the desired concentration and mixed in 1:1 ratio with the different concentrations of SH2 domain (specific fluorescent peptide concentrations can be found in **Table S2**). The mixture was transferred to a black 96-well plate and incubated for 15 minutes at room temperature. Parallel and perpendicular measurements were taken using the 485/30 polarization cube on the BioTek Neo2 Plate Reader. Data was analyzed and fitted to a quadratic binding equation to determine the $K_D$ for the fluorescent peptide, according to previously established methods (2, 3). Next, a peptide of interest was serially diluted 15 times in assay buffer, with the highest concentration being in the high micromolar range (e.g. 400 µM, for a final concentration of 200 µM). In parallel, a fluorescent peptide was mixed with SH2 domain in assay buffer at 2x the desired final concentration (see **Table S2**). The fluorescent peptide/SH2 mixture was mixed 1:1 with the diluted peptides in a black 96-well plate and incubated for 15 minutes at room temperature. Fluorescence polarization was measured as previously described for initial $K_D$ measurements. Competition binding data were fit to a cubic binding equation as described previously (2, 3). P-values comparing affinities for different SH2 domains were calculated using an unpaired Welch's t-test.

### SHP2 activation assays using phosphopeptides

Full-length SHP2 constructs were diluted in assay buffer (60 mM HEPES pH 7.2, 75 mM KCl, 75 mM NaCl,1 mM EDTA, 0.05% Tween-20, with 0.5 mM TCEP freshly added) to a 2X concentration of 0.1 nM. Peptides were serially diluted in assay buffer, with the last point of the concentration series fully without peptide. 15 µL of peptide and 15 µL of 800 µM DiFMUP was added to a black 96-well half area plate. Right before starting the plate reader assay, 30 µL SHP2 was added to each well a final concentration of 0.05nM. Kinetic measurements (Ex 358/Em 355) were taken on a BioTek Neo2 plate reader every 25 seconds for 6 minutes.

### Cloning of the PD-1 ITIM scanning mutagenesis library

A scanning mutagenesis library derived from PD-1 residues 218 to 228 was cloned as described previously for other peptide display libraries in the eCPX system (2, 4). A series of oligonucleotides spanning the peptide-coding sequence was synthesized with a different NNS codon in place of each wild-type codon (11 oligonucleotides in total). These degenerate primers were pooled then used to amplify a library of linear DNA encoding mutant ITIM fusions to the eCPX scaffold protein. This library was then cloned into the pBAD33 vector used for surface display.

### SH2 specificity profiling using bacterial peptide display

*Preparation of bacterial cells*

Electrocompetent MC1061 cells were transformed with ~100 ng of the respective library. After 1 hour recovery in 1 mL LB, cells were further diluted into 250 mL LB + 0.1% chloramphenicol. 1.8 mL of overnight culture was used to inoculate 100 mL LB + 0.1% chloramphenicol, and grown until $OD_{600}$ reached 0.5. 20 mL of cell suspension was induced at 25 °C using a final concentration of 0.4% arabinose until the $OD_{600}$ ~ 1 (after approximately 4 hours). The cells were spun down at 4000 rpm for 15 minutes, and the pellet was resuspended in PBS so that the $OD_{600}$ ~1.5. The cells were stored in the fridge and used within a week.

*Preparing the SH2-beads*

For each sample, 75 µL of Dynabeads™ FlowComp™ Flexi Kit were washed twice in 1 mL SH2 buffer (50 mM HEPES pH 7.5, 150 mM NaCl, 1 mM TCEP, and 0.2% BSA) on a magnetic rack. The beads were then resuspended in 75 µL of SH2 buffer. SH2 domains were thawed quickly and 20 µM of protein was added to the beads. SH2 buffer was added up to 150 µL, and the suspension was incubated for 1 hour at 4 °C while rotating. After 1 hour, the suspension was placed on a magnetic rack and washed twice with 1 mL SH2 buffer.

*Preparing the phosphorylated cells*

150 µL of prepared cells per sample were spun down for 4000 rpm for 5 minutes at 4 °C. Kinase screen buffer was prepared (50 mM Tris, 10 mM magnesium chloride, 150 mM sodium chloride; add 2 mM sodium orthovanadate and 1 mM TCEP fresh) and the cells of each sample were resuspended in 100 µL kinase screen buffer. Kinases c-Src, c-Abl, AncSZ, Eph1B were added to a final concentration of 2.5 µM each, creatine phosphate was added to a final concentration of 5 mM, and phosphokinase was added to a final concentration of 50 µg/mL. The suspension was incubated at 37 °C for 5 minutes before ATP was added to a final concentration of 1 mM. This mixture was incubated at 37 °C for 3 hours. After 3 hours, EDTA was added to a final concentration of 25 mM to quench the reaction. The input library control sample was not phosphorylated. These cells were spun down at 4000 rpm for 15 minutes at 4 °C. The cells were then resuspended in 100 µL of SH2 buffer + 0.1% BSA. Phosphorylation of the cells was confirmed by labeling with the PY20-PerCP-eFluor 710 pan-phosphotyrosine antibody followed by analysis via flow cytometry.

*Enriching for cells displaying high-affinity phosphopeptide ligands*

100 µL of phosphorylated cells were mixed with 75 µL SH2-beads for 1 hour at 4 °C while rotating. After 1 hour, samples were placed on a magnetic rack, supernatant was removed and 1 mL SH2 buffer was added to each sample. This was rotated for 30 minutes at 4 °C to wash the beads. After this wash, the beads were placed on a magnetic rack, the supernatant was removed and 50 µL MilliQ was added.

*Preparing sequencing samples & deep sequencing*

All SH2-selected samples and the input library control were resuspended in 50 µL MilliQ water, vortexed, and boiled for 10 minutes at 100 °C. The boiled lysate was used as the DNA template in a PCR reaction using the TruSeq-eCPX-Fwd and TruSeq-eCPX-Rev primers. The mixture resulting from this PCR was used directly into a second PCR to append Illumina sequencing adaptors and unique 5' and 3' indices to each sample (D700 and D500 series primers). The resulting PCR mixtures were run on a gel, the band of the expected size was extracted and purified, and its concentration was determined using QuantiFluor® dsDNA System (Promega). Samples were pooled at equal molar ratios and sequenced by paired-end Illumina sequencing on a MiSeq or NextSeq instrument using a 150 cycle kit. The number of samples per run, and the loading density on the sequencing chip, were adjusted to obtain at least 1-2 million reads for each index/sample.

*Analysis of deep sequencing data*

Deep sequencing data were processed and analyzed as described previously (2, 4). First, paired-end reads were merged using FLASH (5). Then, adapter sequences and any constant regions of the library flanking the variable peptide-coding region were removed using Cutadapt (6). Finally, these trimmed files were analyze using in-house Python scripts in order to count the abundance of each peptide in the library, as described previously (https://github.com/nshahlab/2022_Li-et-al_peptide-display) (2). The resulting raw counts ($n_{peptide}$) were normalized to the total number of reads in the sample ($n_{total}$) to yield a frequency ($f_{peptide}$) for each peptide in the library (equation 1). The enrichment score for each peptide ($E_{peptide}$) was calculated by taking the ratio of the frequency of that peptide in the enriched sample versus an input (unenriched) sample (equation 2). In the case of the PD-1 ITIM scanning mutagenesis libraries, we report the $log_2$-transformed enrichment of a variant normalized to that for the wild-type ITIM sequence (equation 3).

(1) $f_{peptide} = \frac{n_{peptide}}{n_{total}}$ (2) $E_{peptide} = \frac{f_{peptide,enriched}}{f_{peptide,input}}$ (3) $\Delta E_{variant} = log_2 \frac{E_{variant}}{E_{wild-type}}$

### Cell culture and cell biological experiments

*Cell culture*

All cell lines were incubated at 37°C in a tissue culture incubator with 5% CO2. Human embryonic kidney (HEK) 293 cells were cultured in Dulbecco's Modified Eagle Medium (DMEM) supplemented with 10% Fetal Bovine Serum (FBS) and 1% penicillin/streptomycin. SHP2-deficient Jurkat leukemic cell lines (J.SHP2$^{KO}$) and their counterparts reconstituted with either wild-type SHP2 or T42A or E76K mutants (J.SHP2$^{KO}$-SHP2.WT, J.SHP2$^{KO}$-SHP2.T42A, J.SHP2$^{KO}$-SHP2.E76K) were cultured in RPMI medium supplemented with 5% FBS and 2 mM glutamine. Cells expressing either the wild-type or mutant SHP2 were selected using 2 µg/ml puromycin (Gibco, catalog #A11138-03).

*DNA constructs*

The SHP2 gene was cloned from the pGEX-4TI SHP2 WT plasmid from Ben Neel (Addgene plasmid #8322) (1). The PD-1 gene was cloned from the PD-1-miSFIT-4x plasmid, which was a gift from Tudor Fulga (Addgene plasmid #124678) (7). The mouse Gab1 and Gab2 genes were cloned from the FRB-GFP-Gab2(Y604F/Y634F) and FRB-GFP-Gab1(Y628F/Y660F) plasmids, which were a gift from Andrei Karginov (Addgene plasmid #188658 and #188659) (8). The mouse c-Src gene was expressed from the pCMV5 mouse Src plasmid, a generous gift from Joan Brugge and Peter Howley (Addgene plasmid #13663). The C-terminal regulatory tail (residues 528-535) was deleted to generate hyperactive c-Src. For transient transfection, genes of interest were cloned into a pEF vector (a gift from the Arthur Weiss lab).

*Co-immunoprecipitation experiments*

2.2 x $10^6$ HEK 293 cells were seeded in a 10 cm plate. The next day, cells were transfected using 5 µg of each plasmid (SHP2, Src, interacting protein of interest), and 45 µg PEI in 1.5 mL DMEM. The transfection medium was refreshed after 16 hours and replaced with complete medium. 48 hours later, the cells were harvested by scraping in PBS. Cells were washed 3 times in PBS, and lysed in 500 µL lysis buffer (20 mM Tris-HCl, pH 8.0, 137 mM NaCl, 2 mM EDTA, 10% glycerol, and 0.5% NP-40 + protease inhibitors + phosphatase inhibitors) for 30 minutes while rotating at 4 °C. Cells were spun at 17.7 rpm for 15 minutes at 4 °C. Supernatant was transferred to a clean Eppendorf tube and stored at -20 °C.

Protein concentration was determined using a bicinchoninic acid (BCA) assay and absorbance was measured at 562 nm using a BioTek Synergy Neo2 multi-mode reader. 300 µg of protein in a total volume of 380 µL was incubated overnight with 30 µL of magnetic Myc-beads while rotating at 4

°C. The next day, the beads were washed 3 times on a magnetic rack using 1 mL lysis buffer. Then, 65 uL1x Laemmli buffer was added and beads were boiled at 100ºC for 8 minutes. For whole cell lysates, 15 µg protein was loaded onto a gel. For IP samples, 15 µL of boiled supernatant was used. Gel was transferred onto a nitrocellulose membrane using TurboBlot (BioRad) and the membrane was blocked using 5% bovine serum albumin (BSA) in Tris-buffered saline (TBS) for 1 hour at room temperature. Membranes were rinsed with TBS with 0.1% Tween-20 (TBS-T) and incubated with primary antibodies in TBST + 5% BSA overnight at 4ºC (Src 1:1000, β-actin 1:5000, Myc 1:5000, FLAG 1:5000, PD-1 1:1000, pTyr 1:2000). Co-immunoprecipitation of Gab1/Gab2 was detected using an α-FLAG antibody and PD-1 was detected using a PD-1-specific antibody. Membranes were washed and incubated with secondary antibodies (IRDye 680 and 800). Blots were imaged on a LiCor Odyssey. Band intensities were quantified using Image Studio Lite (Version 5.2) using the Median Background setting. IP intensities were divided by corresponding intensities in total cell lysate. The resulting $SHP2^{T42A}$ intensity was divided by the $SHP2^{WT}$. P-values were calculated in GraphPad Prism (Version 10.1.0) using a paired, one-sided T-test.

*EGF-stimulation experiments*

2.2 x $10^6$ HEK 293 cells were seeded in a two 10 cm plates. The next day, cells were transfected using 5 µg of each plasmid (SHP2, and an interacting protein of interest), and 30 µg PEI in 1 mL DMEM. The transfection medium was refreshed after 16 hours and replaced with serum-free DMEM. 48 hours later, the cells were harvested by scraping in PBS. Cells from both plates were mixed and washed 3 times in PBS. At the third wash, $1/5^{th}$ of the suspension volume was transferred to a separate Eppendorf tube and kept on ice. The remainder of the cell suspension pelleted and resuspended in 1mL of pre-warmed PBS with 25 ng/mL EGF, and placed in a 37°C heat block. At each time point, the cell suspension was mixed by pipetting, and 250 µL was transferred to a new Eppendorf tube. This was immediately centrifuged at 1000g for 5 minutes in at tabletop centrifuge at 4°C. The supernatant was aspirated and the cells were lysed in lysed in 100 µL lysis buffer (20 mM Tris-HCl, pH 8.0, 137 mM NaCl, 2 mM EDTA, 10% glycerol, and 0.5% NP-40 + protease inhibitors + phosphatase inhibitors) for 30 minutes on ice. Cells were spun at 17.7 rpm for 15 minutes at 4 °C. Supernatant was transferred to a clean Eppendorf tube and stored at -20 °C.

Protein concentration was determined using a bicinchoninic acid (BCA) assay and absorbance was measured at 562 nm using a BioTek Synergy Neo2 multi-mode reader. 15 µg protein was loaded onto a gel. Gel was transferred onto a nitrocellulose membrane using TurboBlot (BioRad) and the membrane was blocked using 5% bovine serum albumin (BSA) in Tris-buffered saline (TBS) for 1 hour at room temperature. Membranes were rinsed with TBS with 0.1% Tween-20 (TBS-T) and incubated with primary antibodies in TBST + 5% BSA overnight at 4ºC (Erk 1:2000, p-Erk 1:1000, Vinculin 1:1000, Myc 1:5000, FLAG 1:5000). Membranes were washed and incubated with secondary antibodies (IRDye 680 and 800). Blots were imaged on a LiCor Odyssey. Band intensities were quantified using Image Studio Lite (Version 5.2) using the Median Background setting. p-Erk intensities were divided by total Erk intensities, and normalized to the intensity of $SHP2^{WT}$ at T=2. P-values were calculated in GraphPad Prism (Version 10.1.0) using a paired, one-sided T-test.

*Generation of SHP2-deficient Jurkat cells*

$J.SHP2^{KO}$ cells were generated using Crispr/Cas9 technology as previously described (9, 10). In brief, guide single guide RNAs (sgRNAs) against human SHP2 coding regions were cloned into the pU6-(BbsI)CBh-Cas9-T2A-BFP vector (Addgene, plasmid 64323), and electroporated into Jurkat T cells. The sgRNA sequences used were as follows: 5'-CACCGAGACTTCACACTTTCCGTT-3'; 5'-AAACAACGGAAAGTGTGAAGTCTC-3'. Single clones lacking SHP2 were identified through immunoblot analysis using an anti-SHP2 antibody (clone EPR17829-9; Abcam, catalog # Ab187040). All experiments were done with clone D2 (**Figure S7E**).

*Production of retrovirus expressing wild-type, T42A, or E76K SHP2 mutants*

Human wild-type or mutant SHP2 was cloned into MSCV backbone. The fluorescence protein GFP was incorporated to assess transduction efficiency and expression level. The MSCV plasmids expressing SHP2 variants and packaging envelop plasmids pAmpho were transiently co-transfected into GP2-293 packaging cells (Takara, catalog # 631530) using Xfect Transfection Reagent (Takara, catalog # 631317). Supernatants containing virus particles were collected 48 h after transfection.

*Reconstituted SHP2-deficient Jurkat cells with SHP2 mutants*

J.SHP2$^{KO}$ cells underwent retroviral transduction with viruses expressing SHP2$^{WT}$, SHP2$^{T42A}$, or SHP2$^{E76K}$, by spinning down at room temperature for 1 hour at 2,200 rpm (spinfection), in media supplemented with lipofectamine (Thermofisher, # 11668019). The cells were incubated for 3 days in RPMI medium supplemented with 5% FBS and 2 mM glutamine. Following this, the cells were under drug selection using 2 μg/ml puromycin. The successful reconstitution and expression levels were verified by monitoring GFP expression via LSRFortessa (BD Biosciences).

*CD69 activation assay*

A series of titrated concentration of anti-human CD3 mAb (clone OKT3; Tonbo Biosciences, catalog #70-0037-U100) was used to coat 96-well plates at 4 °C overnight. The next day, the plates were washed once with PBS, and then 4 × $10^5$ cells of J.SHP2$^{KO}$-SHP2.WT, J.SHP2$^{KO}$-SHP2.T42A, or J.SHP2$^{KO}$-SHP2.E76K cells were added to each well. The plates were incubated at 37 °C for 12–16 hours. Cells were stained with anti-CD69 Ab (clone FN50; BD Biosciences, catalog #561928) at 1:800 dilution in FACS buffer (0.5% BSA, 1 mM EDTA in PBS buffer) for 1 hour on ice. Upregulation of CD69 was analyzed on an LSRFortessa (BD Biosciences).

*Phospho-Erk activation assay*

SHP2 variant cell lines were washed in PBS and barcoded with 5-fold dilutions of CellTrace Violet (ThermoFisher #C34557) at 4 × $10^6$ cells/ml for 5 minutes at room temperature. Cells were washed in FACS buffer containing 0.5% BSA in order to deactivate excess dye in solution before being combined and stimulated together. Cells were then stimulated in RPMI at 37 °C with 1:1000 anti-TCR Ab (clone C305) for indicated time. Stimulation was stopped by fixation with 4% Paraformaldehyde (Electron Microscopy Sciences, catalog #15710) for 20 minutes at room temperature before being resuspended in ice cold methanol and kept at 4 °C overnight. Fixed cells were then washed in FACS buffer and stained for two hours at room temperature with unconjugated rabbit anti-pERK antibody (clone 197G2; Cell Signaling Technologies, catalog #4377S) at 1:200 dilution. Next, cells were washed and stained with APC-conjugated goat anti-rabbit IgG antibody (Jackson ImmunoResearch, catalog #711-136-152) for 30 minutes at room temperature. Cells were then washed in FACS buffer and analyzed on an LSRFortessa (BD Biosciences).

### Melting curves

Purified protein stocks were thawed and diluted in DSF buffer (20 mM HEPES pH 7.5, 50 mM NaCl, 0.4% DMSO. 19 µL was added to a MicroAmp Fast Optical 96-well Reaction plate (Applied Biosystems, # 4346906). 1 µL of 500x SYPRO Orange Protein Gel Stain (Thermo Fisher, catalog no. S-6650) was added to a final protein concentration of 10 µM and 25x SYPRO Orange. Melting curves were performed in an Applied Biosystems Step-One Plus RT-PCR thermocycler. Temperature measurements started at 15 ºC or 25 ºC, and temperature was raised by 0.5 ºC every minute with continuous measurements of fluorescence (excitation: 472 nm; emission: 570 nm). Raw fluorescence values along with corresponding temperatures were analyzed using DSFworld (11) and $T_m$ values were calculated using dRFU.

### Molecular Dynamics Simulations

*Preparation of Structural Models for Simulations*

We built and simulated several systems comprising of the SHP2 N-terminal SH2 domains bound to different ligands, as well as the SH2 domain in the apo state. For most of the systems, the SH2 domain was taken from the crystal structure 6ROY (12). This structure, without a ligand bound, was used as the starting structure for simulations of the SH2 domain in the apo state. For simulations of the SH2 domain bound to PD-1 pTyr$^{223}$, the ligand was taken from the PDB structure of 6ROY, and the missing residues -6 (Pro), -5 (Val), -4 (Phe) and -3 (Ser) were built in using PyMOL (13). For simulations of the SH2 domain bound to IRS-1 pTyr$^{896}$, the structures of the SH2 domain and the bound ligand were taken from the PDB structure of 1AYB (14). Here the N-terminal residues 1-4 of the SH2 domain were mutated to MTSR (from MRRW) to be consistent with the sequence of the SH2 domain in the rest of the simulations. For the same reason, Cys was built in at position 104 at the C-terminal end. Residues -6 (Phe), -5 (Lys), -4 (Ser), and -3 (Pro) in the ligand were missing in the crystal structure and were built in using PyMOL. For simulations of the SH2 domain bound to Gab2 $pTyr^{614}$, the structure of the SH2 domain and ligand were taken from the PDB structure of 6ROY. The starting structure of the SH2 domain was built in the same way as that used in the simulations of the SH2 domain bound to PD-1. The ligand was built by mutating residues -6, -5, -4, +1, and +2 of the ligand in simulations of PD-1 bound to the SH2 domain to Ser, Thr, Gly, Leu, and Ala, respectively. For the simulations of the SH2 domain bound to Imhof-9, the PDB structure 3TL0 was used (15). For the starting structure of the SH2 domain, missing residues 1-4 (MTSR) at the N-terminal end and residue 104 (Cys) at the C-terminal end were built in using PyMOL. For the starting structure of the ligand, missing residues -6 to -3 (KKAA) were built in, residue +4 was mutated from Tyr to Leu and missing residues +4 to +6 (MFP) were built in using PyMOL. For the simulations of the SH2 domain bound to MILR1 $pTyr^{338}$, the crystal structure from 6ROY was used. The SH2 domain was built in the same manner as in simulations of the SH2 domain bound to PD-1. For the ligand, residues -6 to -1 of the ligand in 6ROY were mutated to AKSGAV (from PVFSVD), residue +1 was mutated from Gly to Ser, residue +4 was mutated from Asp to Asn, and residue +6 was mutated from Gln to Gly. For each of these systems, a similar system with $Thr^{42}$ in the SH2 domain mutated to alanine was built using PyMOL. Crystalline waters from 6ROY were used in all the simulations. In all the systems, the N-terminal and C-terminal ends were capped with acetyl and amide groups respectively, in both the SH2 domains and the ligands.

*Simulation Protocol*

All the systems were solvated with TIP3P water (16) and ions were added such that the final ionic strength of the system was 100 mM using the tleap package in AmberTools21 (17). The energy of each system was minimized first for 5000 steps while holding the protein chains and crystalline waters fixed, followed by minimization for 5000 steps while allowing all the atoms to move. For each system, three individual trajectories were generated by reinitializing the velocities at the start of the heating stage described below.

The temperature of each system was raised in two stages – first to 100 K over 1 ns and then to 300 K over 1 ns. The protein chains and crystalline waters were held fixed during this heating stage. Each system was then equilibrated for 2 ns, followed by production runs. Three production trajectories, each 1 μs long, were generated for each system. All equilibration runs and production runs were performed at constant temperature (300 K) and pressure (1 bar).

The simulations were carried out with the Amber package (18) using the ff14SB force field for proteins (19) using an integration timestep of 2 fs. The Particle Mesh Ewald approximation was used to calculate long-range electrostatic energies (20). All hydrogens bonded to heavy atoms were constrained with the SHAKE algorithm (21). The Langevin thermostat was used to control the temperature with a collision frequency of 1 $ps^{-1}$. Pressure was controlled while maintaining periodic boundary conditions.

*Analyses*

Key measurements were extracted from the MD trajectories using the CPPTRAJ module of AmberTools22 (22). For the RMSF calculations, the trajectories were sampled every 100 ps and RMSF values were calculated from the Cα atoms of each residue after determining root mean squared deviation relative to the first state in the production run of the simulation. For RMSF calculations of each system, each trajectory was analyzed separately, as seen in Figure S5A. For distance calculations, trajectories were sampled every 1 ns. The distance measurements from all three replicates of each system were combined to determine the distance distributions seen in Figure 5, Figure S5, and Figure S6. In cases where distance calculations involved a redundant atom (e.g. distances to one of the three non-bridging oxygen atoms in the phosphoryl group of phosphotyrosine), all three distance measurements were calculated, then the shortest distance at each frame was determined and used for the distribution plots. For visualization, trajectories were sampled every 10 ns. All structure visualization and rendering in this study was done using PyMOL (13).

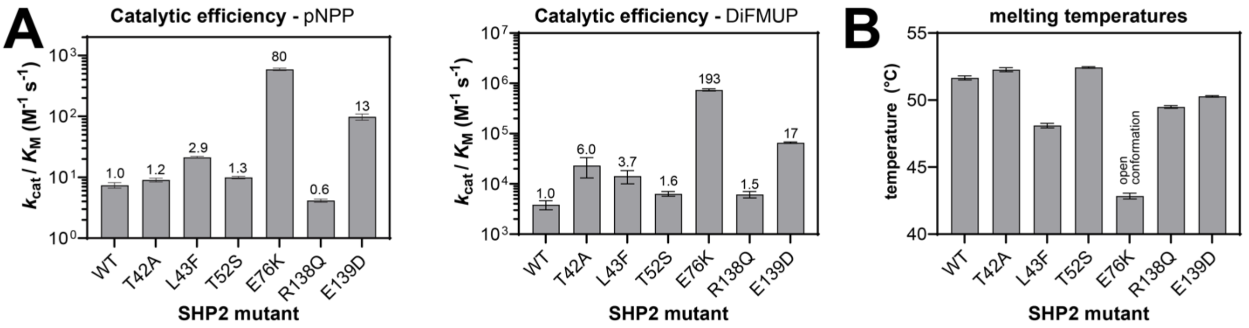


**Figure S1. Basal activity measurements and melting temperatures of various SHP2 mutants.** (**A**) Left panel: Catalytic efficiency of SHP2 mutants against *p*-nitrophenyl phosphate (pNPP). N = 3 separate dilutions of protein and titrations of pNPP stock. Right panel: catalytic efficiency of SHP2 mutants against 6,8-difluoro-4-methylumbelliferyl phosphate (DiFMUP). N = 3-6 separate dilutions of protein and titrations of DiFMUP stock Fold-change compared to wild-type is indicated on top of each bar. Numerical values for catalytic efficiency can be found in Table S1. (**B**) Melting temperatures of SHP2 mutants. N = 3-9 separate dilutions of protein. Numerical values for melting temperatures can be found in Table S1.

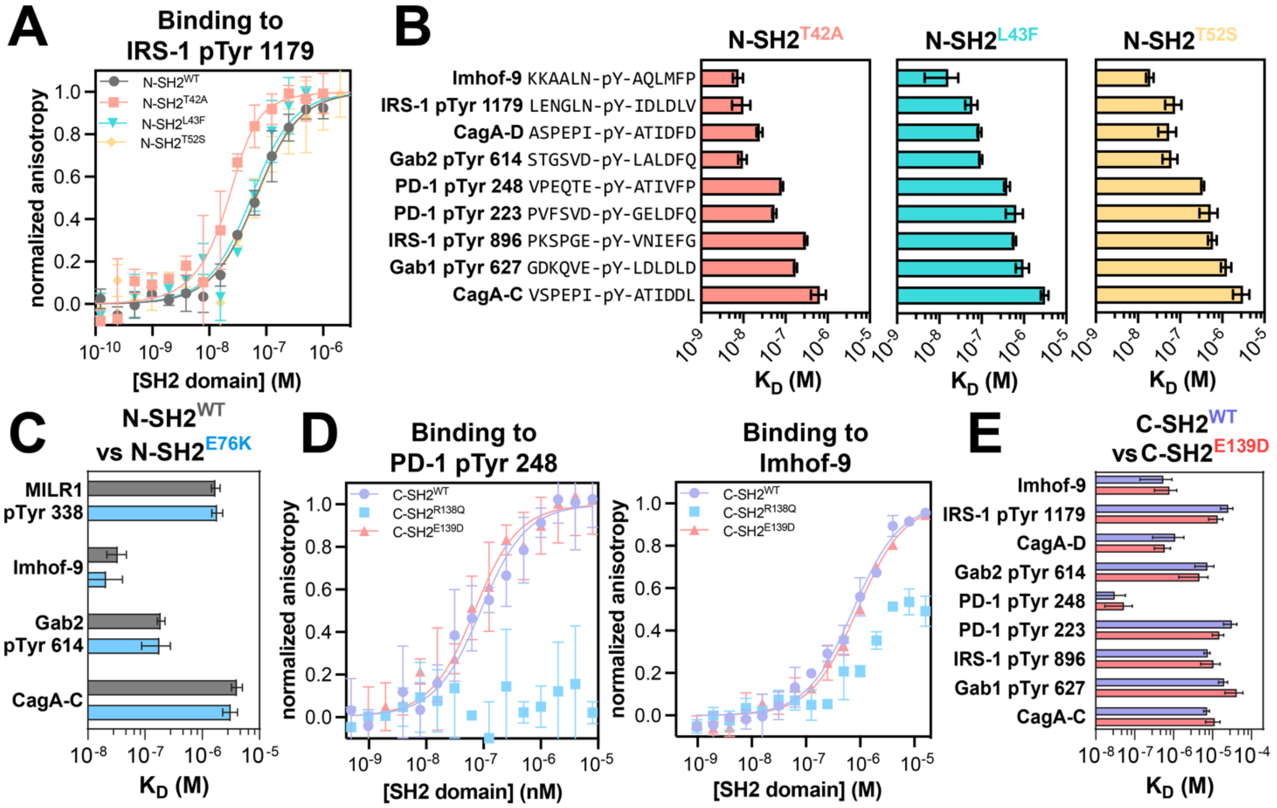


**Figure S2. Binding affinities for SH2 domains.** (**A**) Fluorescence polarization binding measurements of N-SH2 domains against the fluorescently-labeled IRS-1 pTyr 1179 peptide. N = 3-9 independent protein and fluorescent peptide dilutions. (**B**) Binding affinities of N-SH2$^{T42A}$, N-SH2$^{L43F}$, and N-SH2$^{T52S}$ for various peptides derived from known SHP2 interactors. N = 3-5 independent protein, peptide, and fluorescent peptide titrations. (**C**) Binding affinities of N-SH2$^{WT}$ and N-SH2$^{E76K}$ against select peptides derived from known SHP2 interactors. N = 3-4 independent protein, peptide and fluorescent peptide titrations. (**D**) Binding affinity measurements for C-SH2$^{WT}$, C-SH2$^{R138Q}$, and C-SH2$^{E139D}$ against fluorescently labeled PD-1 pTyr 248 and Imhof-9 phosphopeptides. N = 3-6 independent protein, and fluorescent peptide titrations. (**E**) Binding affinities of C-SH2$^{WT}$ and C-SH2$^{E139D}$ with various phosphopeptides derived from known SHP2 interactors. N = 3-6 independent protein, peptide, and fluorescent peptide dilutions. Source data for binding affinities can be found in Table S2.

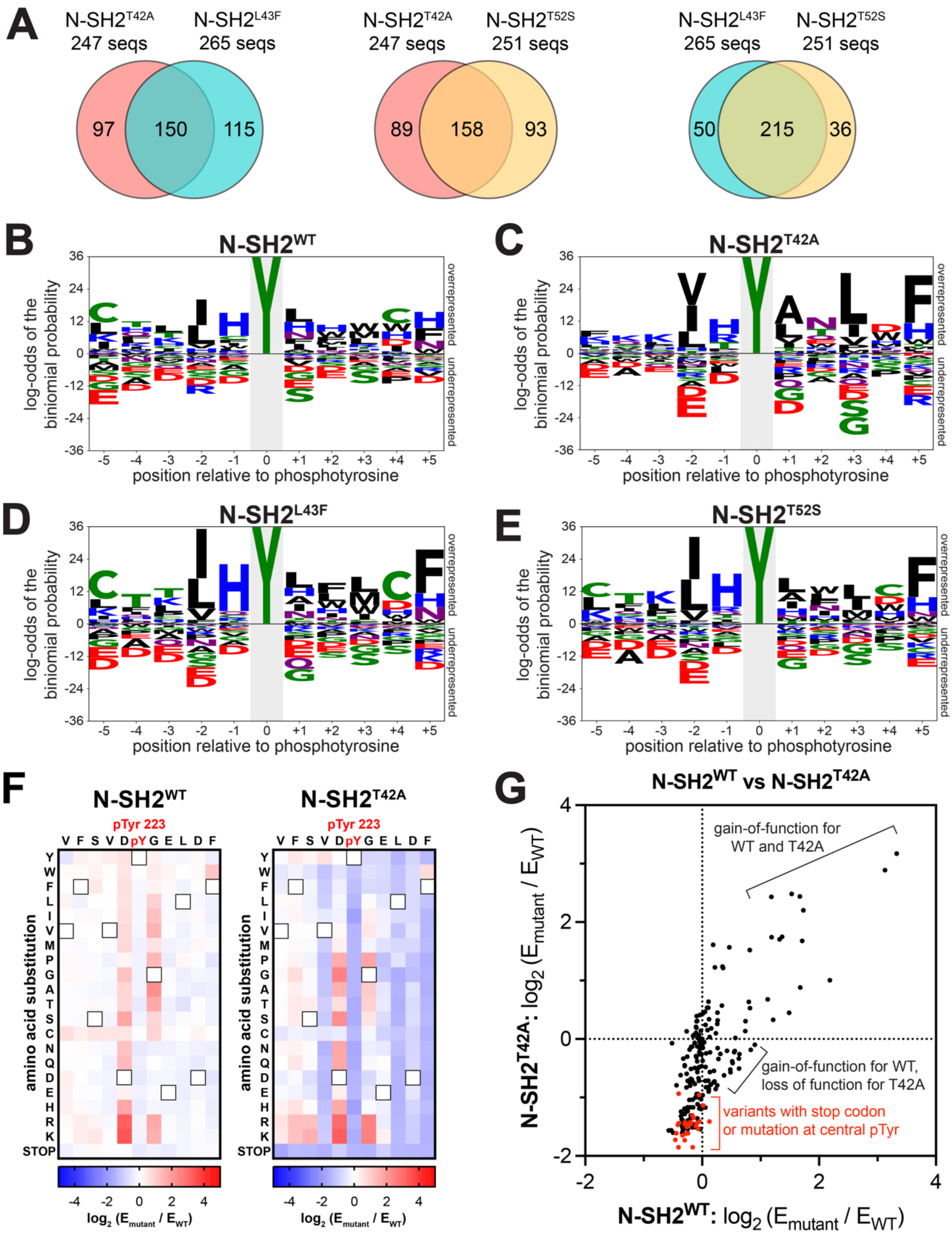


**Figure S3. Analysis of sequence features for N-SH2 mutants.** (**A**) Overlap in sequences enriched by each mutant N-SH2 domain above an enrichment score cutoff of 3.2. (**B**) Probability logo for N-SH2$^{WT}$, indicating the probability at each position of the amino acids in the peptides with an enrichment score above cut-off of 3.2 (168 peptides) relative to all peptides in both libraries (9281 peptides). (**C**) Probability logo, as in (B), for N-SH2$^{T42A}$ (247 peptides). (**D**) Probability logo, as in (B), for N-SH2$^{L43F}$ (265 peptides). (**E**) Probability logo, as in (B), for N-SH2$^{T52S}$ (251 peptides). (**F**) Heatmaps of N-SH2$^{WT}$ and N-SH2$^{T42A}$ for PD-1 pTyr$^{223}$ (ITIM) scanning mutagenesis screen (N = 5 independent library transformations and screens). The wild-type residue on each position is indicated by a black square. Color scheme: blue indicates weaker binding, white indicates same as wild-type, red indicates tighter binding. Source data for the heatmaps in panel (F) can be found in Table S4. (**G**) Correlation plot of the substitution scores for N-SH2$^{WT}$ and N-SH2$^{T42A}$ in the PD-1 pTyr$^{223}$ scanning mutagenesis screens. Negative control substitutions (central pTyr mutants and variants with a stop codon) are shown as red dots.

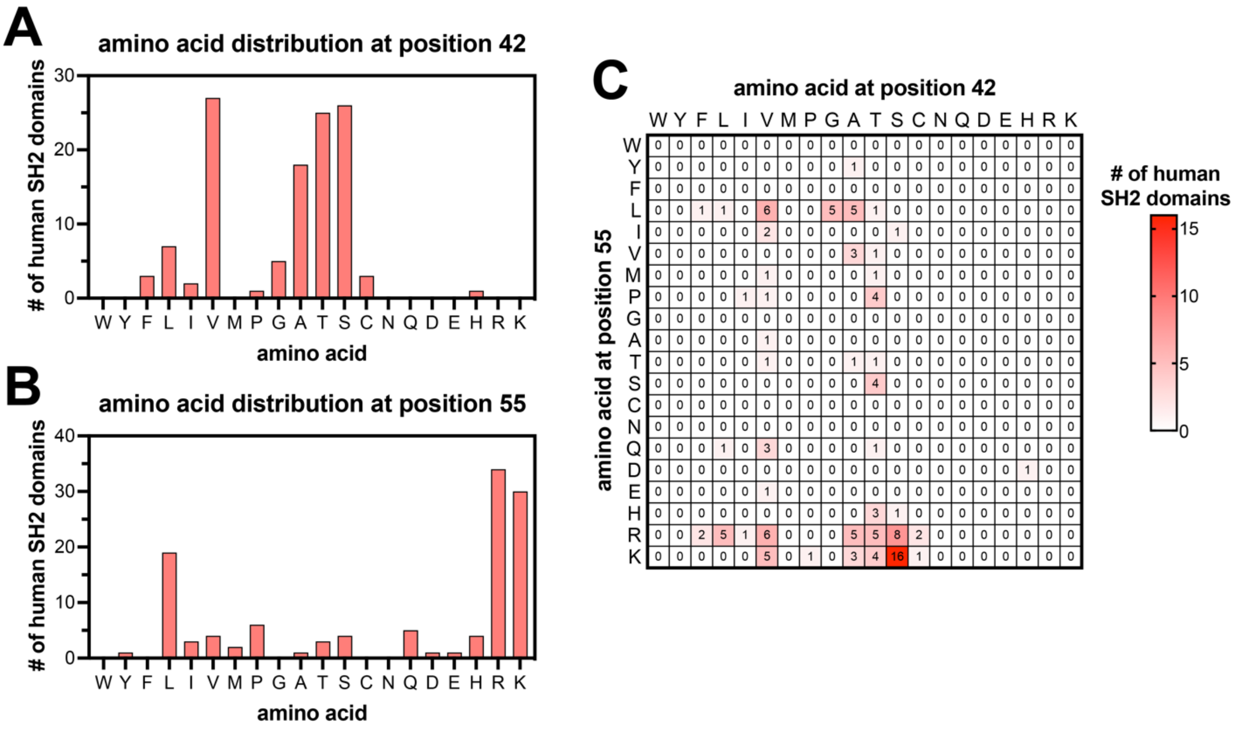


**Figure S4. Amino acid residues at key positions in human SH2 domains.** (**A**) Abundance of each amino acid in 120 human SH2 domains at the position corresponding to $Thr^{42}$ in SHP2 $N\text{-}SH2^{WT}$. (**B**) Abundance of each amino acid in 120 human SH2 domains at the position corresponding to $Lys^{55}$ in SHP2 $N\text{-}SH2^{WT}$. (**C**) Abundance of each pairwise combination of amino acids in the 120 human SH2 domains at positions corresponding to $Thr^{42}$ and $Lys^{55}$ in SHP2 $N\text{-}SH2^{WT}$.

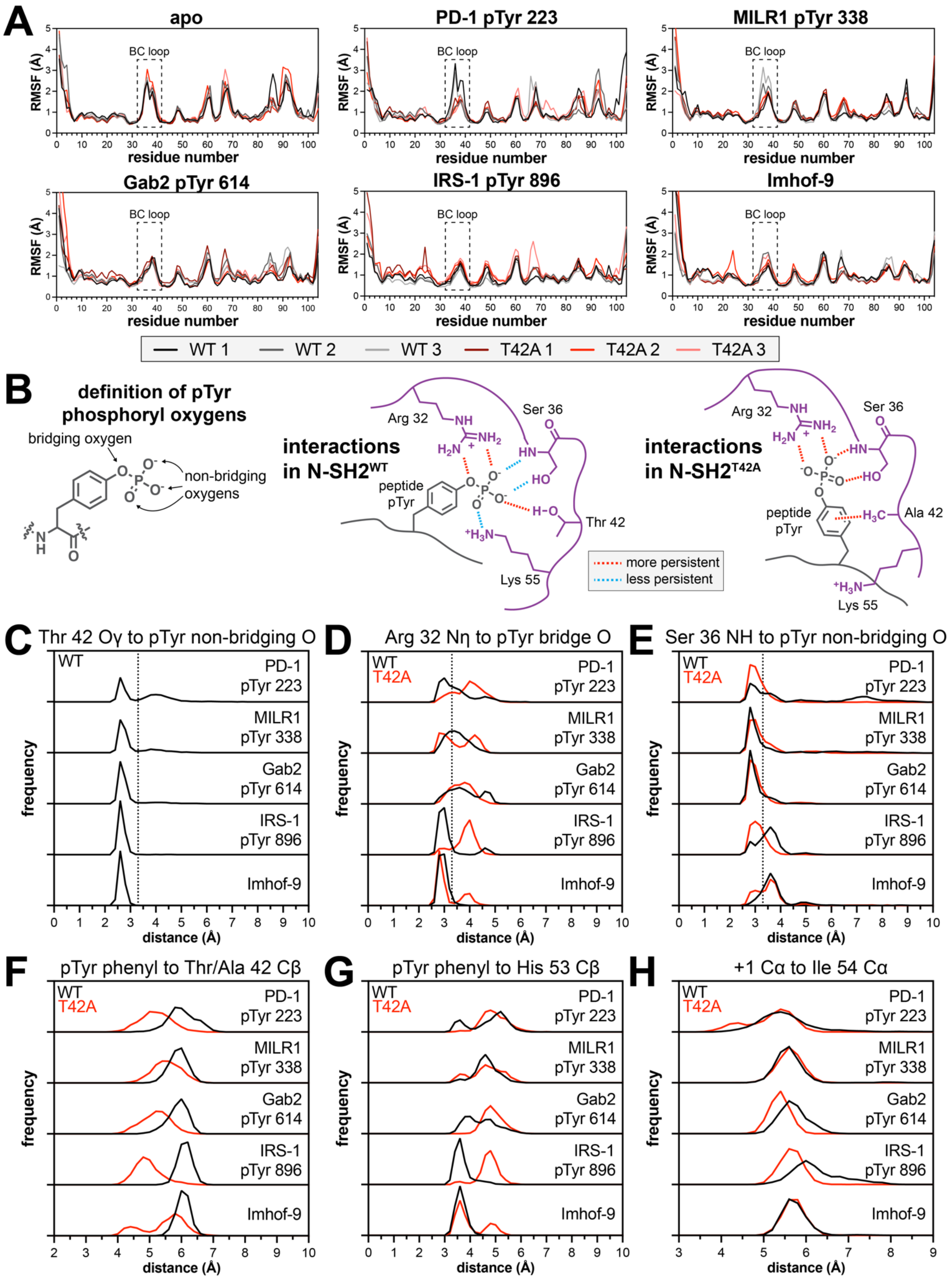


**Figure S5. Measured structural parameters from MD simulations of SHP2 N-SH2$^{WT}$ or N-SH2$^{T42A}$.** (**A**) Root mean squared fluctuation measurements from MD simulations of SHP2 N-SH2$^{WT}$ or N-SH2$^{T42A}$. Calculations were carried out exclusively on Cα atoms and are derived from 1 µs of simulation time, sampled every 100 ps. Peptide-bound simulations show rigidification of the BC-loop. Differences in RMSF values between N-SH2$^{WT}$ and N-SH2$^{T42A}$ were largely negligible in both the apo and peptide-bound states, with one noteworthy exception:

the BC-loop in two-thirds of the N-SH2$^{WT}$ simulations with the PD-1 and MILR1 peptides showed more fluctuation than in the N-SH2$^{T42A}$ simulations. Simulations also show reduced fluctuations around the EF- and BG-loops (residues 64-70 and 86-96, respectively), in the peptide-bound state. These regions recognize the strongly preferred +5 Phe found in all five peptides. (**B**) Two-dimensional illustration of key interactions between the phosphotyrosine residue and the SH2 domain, based on the wild-type and T42A simulations. (**C**) Distance between the Thr$^{42}$ side-chain hydroxyl group and the closest non-bridging phosphoryl oxygen atom. This hydrogen bond broke intermittently in N-SH2$^{WT}$ simulations with the PD-1, MILR1, and Gab2 peptides, but not the tighter binding IRS-1 and Imhof-9 peptides. (**D**) Distance between the pTyr bridging phosphoryl oxygen and the closest Nη atom on Arg$^{32}$. In the N-SH2$^{WT}$ simulations, the bidentate guanidium-phosphoryl interaction involved one non-bridging oxygen and the less electron-rich bridging oxygen, but in the N-SH2$^{T42A}$ simulations, Arg$^{32}$ coordinated two non-bridging oxygens, in a presumably stronger interaction. (**E**) Distance between the amide nitrogen of Ser$^{36}$ and the closest non-bridging phosphoryl oxygen atom. Dotted lines denote an H-bond threshold of 3.3 Å. (**F**) Distance from the pTyr phenyl ring to the Cβ atom of Thr$^{42}$ or Ala$^{42}$. (**G**) Distance from the pTyr phenyl ring to the Cβ atom of His$^{53}$, which lines the peptide binding pocket and plays a role in recognizing the -2 and -1 residues. The phosphotyrosine residue in N-SH2$^{T42A}$ simulations moved further away from His$^{53}$. (**H**) Distance from the Cα atom of Ile$^{54}$ to the Cα atom of the +1 residue on the peptide. All distance distributions are derived from calculations over 3 μs of simulation time (three 1 μs simulations), sampled every 1 ns.

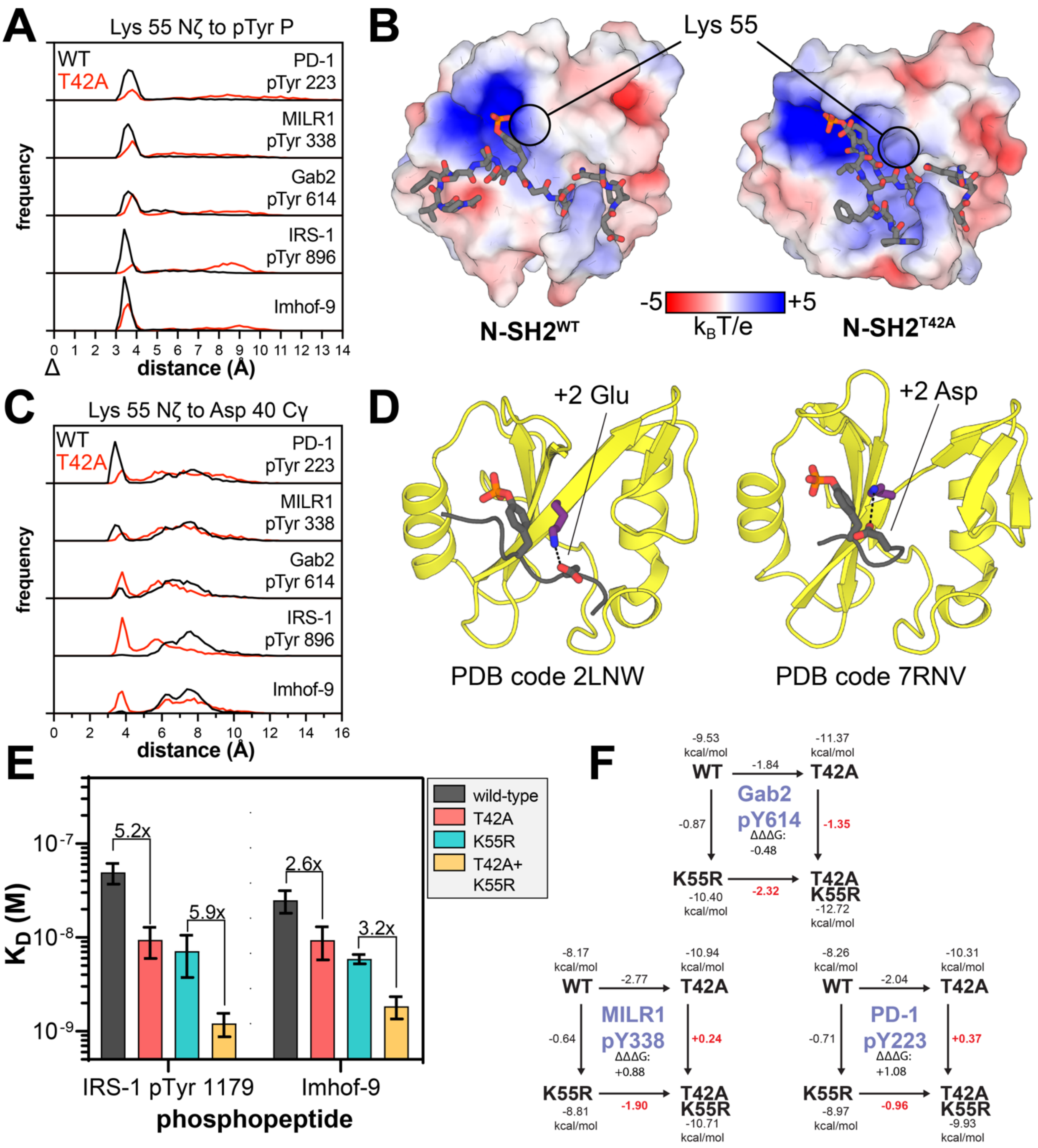


**Figure S6. The role of Lys$^{55}$ on T42A-dependent peptide recognition.** (**A**) Distribution of distances between the Lys$^{55}$ Nζ atom and the phosphotyrosine phosphorus atom in simulations of various peptides bound to N-SH2$^{WT}$ (black) or N-SH2$^{T42A}$ (red). (**B**) Electrostatic surface potential of SHP2 N-SH2$^{WT}$ or N-SH2$^{T42A}$ from select frames in simulations with the PD-1 pTyr$^{223}$ peptide. Surface potentials were calculated using the APBS plugin for PyMOL. The location of Lys$^{55}$ is circled to show how the T42A mutation alters the surface accessibility of Lys$^{55}$. (**C**) Distribution of distances between the Lys$^{55}$ Nζ atom and the Asp$^{40}$ Cγ atom in simulations of various peptides bound to N-SH2$^{WT}$ (black) or N-SH2$^{T42A}$ (red). (**D**) Vav2 SH2 domain structures highlighting interactions with the Lys$^{55}$-analogous lysine residue and acidic residues on the peptide ligand (PDB codes 2LNW and 7RNV). (**E**) Effects of the K55R mutation on peptides for which we observed a small enhancement of binding affinity in context of the T42A mutation. These measurements were made using fluorescently-labeled peptides and direct fluorescence polarization binding measurements, as in Figure S2A. (**F**) Double mutant cycle analysis of Gab2 pTyr 614, MILR1 pTyr 338 and PD-1 pTyr 223 peptides using the T42A and K55R mutants. The difference in sign for ΔΔΔG with Gab2 pTyr 614 when compared to MILR1 pTyr 338 and PD-1 pTyr 223 suggest a different nature of coupling between these residues when a +2 Glu is present.

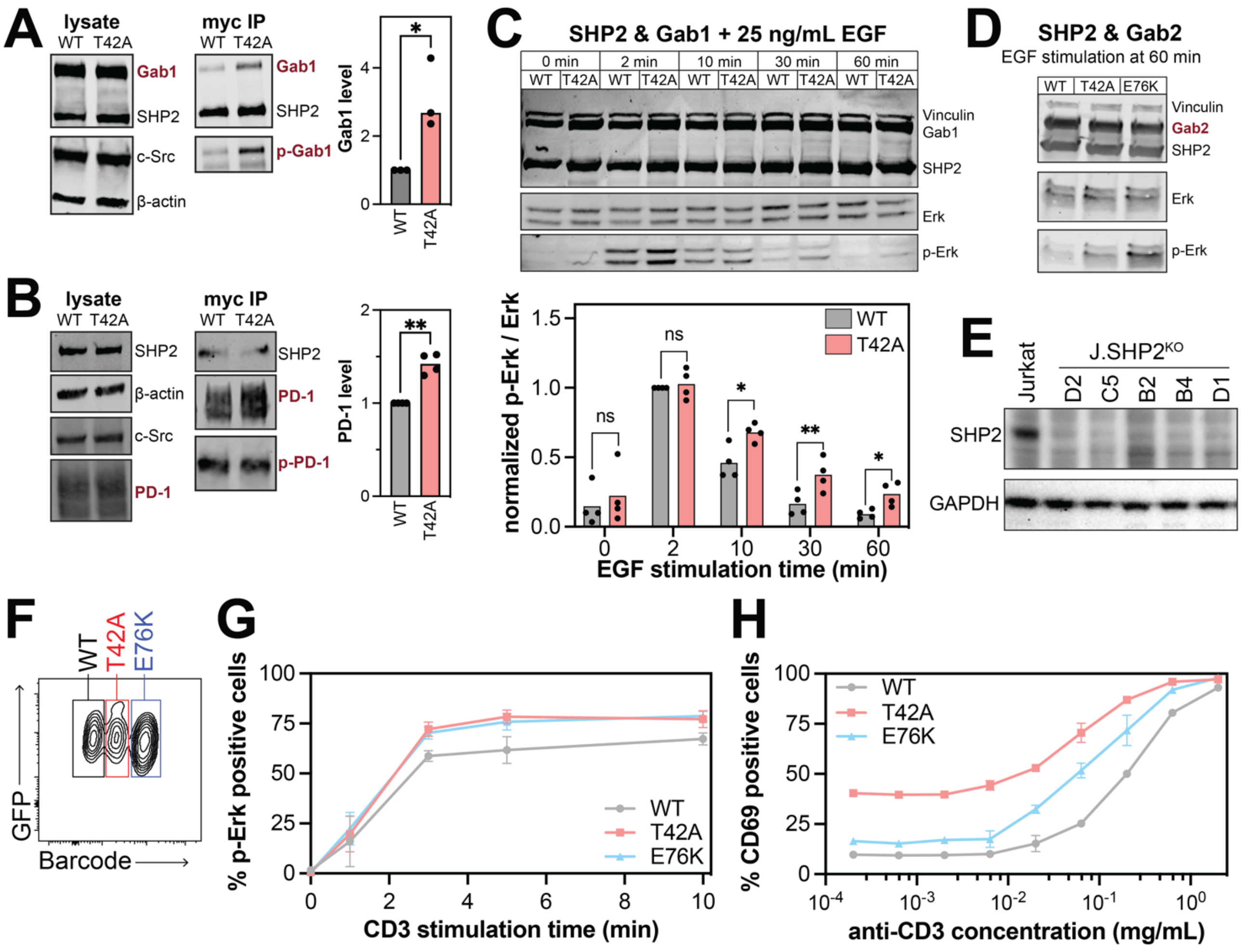


**Figure S7. Cellular consequences of the T42A mutation.** (**A**) SHP2 co-immunoprecipitation results with Gab1 (N = 3). (**A**) SHP2 co-immunoprecipitation results with PD-1 (N = 4). Co-immunoprecipitation levels of each protein in T42A samples relative to wild-type are normalized for expression level and shown as bar graphs. (**C**) phospho-Erk (p-Erk) levels after EGF stimulation in the presence of co-expressed Gab1 and either SHP2$^{WT}$ or SHP2$^{T42A}$ (N = 4). The bar graphs below the blots indicate p-Erk levels, normalized to total Erk levels, relative to the highest p-Erk signal in the SHP2$^{WT}$ time course (2 minutes). For all bar graphs, a paired, one-tailed t-test was used to test for significance. * denotes $p < 0.05$, ** denotes $p < 0.01$. (**D**) Phospho-Erk levels at 60 minutes after EGF stimulation in presence of co-expressed Gab2, related to the experiments shown in Figure 7B. The 60 minute timepoint was not quantified as it was run on a different gel. (**E**) Western blot validation of Jurkat SHP2 knock-out (KO) cell line. All subsequent experiments were done with clone D2. (**F**) SHP2$^{WT}$, SHP2$^{T42A}$ or SHP2$^{E76K}$-reconstituted SHP2 knock-out Jurkat cells were individually labeled with CellTrace Violet dye at different concentrations, then pooled together for the experiments. Representative flow cytometry plot of SHP2 variants-expressing cells barcoded with titrated amounts of CellTrace Violet dye is shown. GFP expression is used as an indicator of SHP2 variant expression level. (**G**) Representative analysis of the pERK+ population in SHP2$^{WT}$, SHP2$^{T42A}$, or SHP2$^{E76K}$-reconstituted SHP2 knock-out Jurkat cells (n =3 replicates of independent stimulation). (**H**) Percentages of CD69+ cells as a function of anti-CD3 antibody concentrations, for SHP2 knock-out Jurkat cells reconstituted with SHP2$^{WT}$, SHP2$^{T42A}$, or SHP2$^{E76K}$ (N = 2 stimulation replicates).
